# An African trypanosome surface protein inhibits amplification of the complement system

**DOI:** 10.64898/2026.08.17.745214

**Authors:** Alexander D. Cook, Helena Webb, Nicola Minshal, Mark Carrington, Matthew K. Higgins

## Abstract

African trypanosomes replicate within the blood and tissue spaces of their mammalian hosts. As extracellular pathogens, they are constantly exposed to immune cells and molecules and are susceptible to killing by components of the complement system. In this study, we conducted a large-scale screen of a panel of human complement components against a set of putative *T. brucei* surface proteins and discovered a novel *T. brucei* receptor that binds to complement Factor B. Biochemical and structural characterisation of this protein revealed it to be a potent inhibitor of the C3bBb convertase, a central enzyme in amplification of the complement cascade. Structural studies show that this C3bBb receptor bridges C3b and Bb in a conformation which is incompatible with its catalytic activity, directly blocking C3bBb convertase function. This reveals a novel mechanism of C3bBb convertase regulation and deepens insight into how the African trypanosome cell surface has evolved to evade complement-mediated killing.

## Introduction

The complement system is a central part of innate and acquired immunity and presents a significant barrier to successful infection by pathogens. It consists of a central proteolytic cascade (Figure 1a), which can be triggered through different mechanisms, resulting in a series of downstream effector functions. Recruitment of complement follows from three pathways: the lectin pathway involving pattern recognition of specific sugars on pathogen surfaces, the classical pathway involving pattern recognition by pentraxins or antibody binding to pathogen antigens, or the alternative pathway involving spontaneous complement activation. In each case, initiation triggers the central proteolytic cascade of complement, resulting in deposition of complement factor C3b on the pathogen surface and formation of the C3bBb convertase which catalyses a positive feedback loop of further C3 deposition. This leads to release of immunostimulatory anaphylatoxins which mediate inflammation and chemotaxis. In addition, deposited surface complement components recruit immune cells, enhance phagocytosis of pathogens^1^ and trigger B cell activation^2^. They also recruit components of the membrane attack complex, resulting in pathogen lysis. Indeed, individuals with complement deficiencies experience recurrent severe infections, particularly of extracellular bacteria, indicating the importance of complement to immunity^3^.

**Figure 1.**
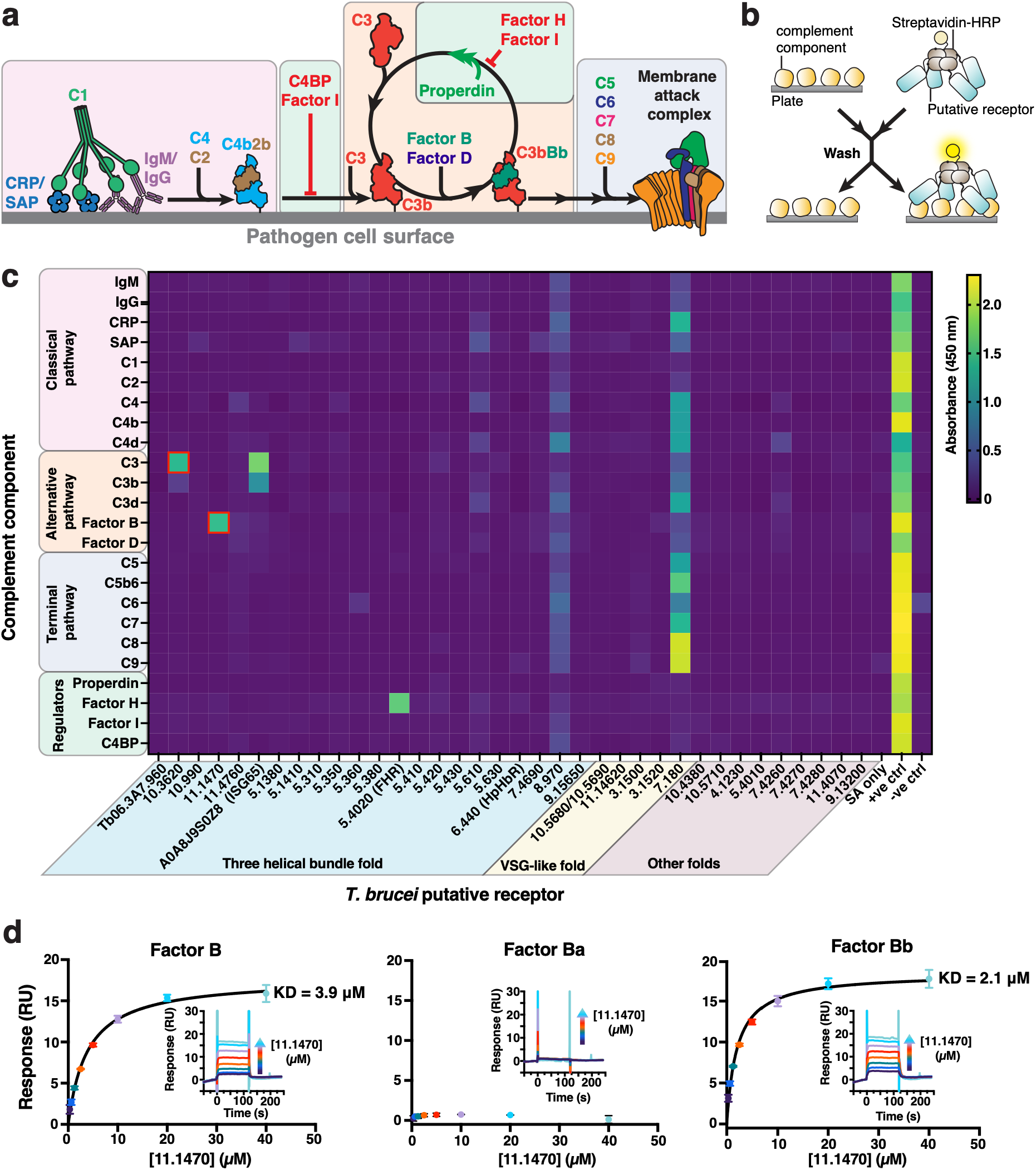
A protein-binding interaction screen shows that Tb927.11.1470 bind to complement Factor. **B.** (a) The core proteolytic cascade of the complement system, highlighting all complement proteins included in the screen. (b) A schematic illustration of the interaction screen format, showing that complement components are non-specifically adsorbed on a 96-well plate, followed by addition of streptavidin-HRP tetramerised *T. brucei* putative receptors. Tetramerised receptors not removed by washes generate a yellow signal. (c) Heatmap showing interactions between streptavidin-HRP tetramerised *T. brucei* putative receptors and immobilised complement components. Red boxes highlight previously unidentified interactions. Absorbance values are an average value from technical replicates, n = 2. (d) Surface plasmon resonance data showing responses from injection of Tb927.11.1470 (two-fold serial dilutions from a concentration of 40 μM) over a flow cell coupled with either complement Factor B, Factor Ba, or Factor Bb. Data is representative of three experiment repeats, standard error of the mean is plotted.

In response, numerous pathogens and biting arthropods have evolved complement binding proteins which inhibit different parts of the cascade. Bacteria (e.g. *Streptococcus^4,5^, Borrelia^6,7^* and *Neisseria* species^8^) and eukaryotic pathogens (e.g. African trypanosomes^9^ and Plasmodium parasites^10^) produce surface molecules which recruit the negative complement regulators Factor H and C4 binding protein, while viral proteins have evolved to mimic Factor H^11^. These regulators block progress of the complement cascade. Sandflies^12^ and leeches^13^ produce protease inhibitors which block initiation of complement through the classical pathway, while ticks produce proteins that inhibit activation of complement factor C5^14^, preventing production of the most potent anaphylatoxin C5a and formation of the membrane attack complex. However, one of the most universal routes to block the complement cascade is to directly inhibit the formation or function of the central C3bBb convertase, with sandfly saliva component Lufaxin preventing formation of C3bBb^15^ and *S. aureus* protein SCIN allowing formation of stable but inactive C3 convertases^16^.

African trypanosomes have been used as a model for how pathogens resist complement-mediated killing. These single-celled parasites live freely in the bloodstream and tissues of their mammalian hosts and are therefore constantly exposed to immune cells and molecules, including complement. This has driven an unusual adaption of their glycoprotein cell surface, with each cell decorated with ∼1 x 10^7^ copies of variant surface glycoprotein (VSG). Antigenic variation, driven by changing which VSG is expressed from thousands of genes, allows evasion of adaptive immunity at the population level^17^. In addition, the African trypanosome surface contains tens of putative receptor proteins^18,19^ with three-helical bundle^20^ or VSG-like architectures^21^. Two of these have been shown to bind complement factors: a Factor H receptor^9^ and the C3 receptor ISG65^22,23^. We therefore asked whether the parasite expresses additional receptors which enable it to evade killing by the complement system.

## Results

### Discovery of a Trypanosoma brucei receptor for Complement Factor B

To identify novel complement receptors, we developed a protein-based screen. We first constructed a library of putative *T. b. brucei* cell surface receptors, starting by performing a bioinformatics search to identify a set of proteins that were predicted to contain a GPI-anchor or a single-pass transmembrane helix. Many trypanosome surface proteins adopt a VSG-like fold or a three helical bundle fold and so we also searched for these protein classes. This yielded a panel of 37 putative surface proteins, corresponding to single genes or representative members of gene families and we classified these based on whether they are VSG-like, three-helical bundles or neither (Supplementary Data 1). All of these proteins were predicted to contain a signal peptide for Sec61/SPI-mediated secretion, with 16 predicted to be type-I transmembrane, 18 to be GPI-anchored, and 3 to be secreted. Tb927.10.5680 and Tb927.10.5690 were predicted to form a heterodimer and were co-expressed.

Next, we screened the panel of putative receptors for binding to a panel of complement components, using an approach based on the AVEXIS method^24^. Putative receptors were recombinantly expressed in HEK293F or S2 cells with a C-terminal biotin replacing the GPI anchor site or transmembrane helix (Supplementary Table 1, Supplementary Figure 1 a), allowing presentation of the receptor on a streptavidin horseradish-peroxidase fusion (Figure 1b). This tetramerises putative receptors and presents them with an orientation matching that on the trypanosome surface, with avidity increasing the sensitivity of the assay.

We also selected a panel of components of the complement system, including components of the core cascade, as well as those specific for the classical and alternative pathways, and those that form the membrane attack complex of the terminal pathway. We included some component regulators as a previous trypanosome receptor has been identified against the regulator Factor H^9^ (Figure 1a). Complement components were either purchased, recombinantly expressed or endogenous purified (Supplementary Figure 1b). Each complement component was immobilised non-specifically on a plate, then each putative multimerised receptor was added to test for binding (Figure 1b). This method resulted in good signal to noise ratio, as evidenced by binding of ISG65 to complement C3 with low background signal (Supplementary Figure 2). Of the 37 *T. b. brucei* proteins expressed, we detected clear binding for the known C3b-binding ISG65 receptor^22,23^ and Factor H receptor^9^. In addition, the screen indicated binding partners for two proteins with no previously known function (Figure 1C, Supplementary Figure 2). These were Tb927.10.3620, which appeared to bind both C3 and C3b, and Tb927.11.1470, which bound to Factor B.

To validate these findings, we used surface plasmon resonance (SPR). We did not observe binding of Tb927.10.3620 to immobilised C3, C3b, and C3d on a surface plasmon resonance chip (Supplementary Figure 3), suggesting that this may be a false positive. In contrast, when we flowed Tb927.11.1470 over immobilised Factor B and its activation products Factor Ba and Factor Bb, we observed clear binding, with affinities of 4 μM for Factor B and 2 μM for Factor Bb, but no binding to Factor Ba (Figure 1d, Supplementary Figure 4, Supplementary Table 2). This confirms that Tb927.11.1470 binds Factor B, most likely interacting with the serine protease domain and/or the von Willebrand domains also found in Factor Bb.

### Tb927.11.1470 inhibits complement amplification

We next determined whether Tb927.11.1470 modulates the activity of Factor B. The sole known function of Factor B is to assemble with C3b to form the alternative pathway C3 convertase to amplify the complement cascade (Figure 1a). To achieve this, Factor B binds to C3b, forming the pro-convertase C3bB. C3bB is recognised by Factor D; a serine protease that cleaves Factor B into Ba and Bb fragments. Factor Bb remains bound to form the active C3bBb convertase, which cleaves additional C3 molecules, increasing the pool of C3b available to form further C3bBb convertases. We therefore asked whether Tb927.11.1470 affects C3bBb convertase mediated amplification.

We first established an assay for C3bBb amplification. We combined 6.5 μM C3, 2.4 μM Factor B and 70 nM Factor D, matching the average concentrations of these proteins in human blood, and added 70 nM C3b to initiate convertase formation (Figure 2a). Initially, a small number of C3bBb convertases form from the C3b provided. We then used SDS-PAGE to measure the relative abundance of Factor Ba and C3a cleavage products resulting from the amplification reaction over time (Figure 2b). Within 5 seconds, Factor Ba and C3a were visible. After calculating relative band density, we observed that C3a and Factor Ba increase steadily between 5 and 20 seconds, before levelling off as C3 and Factor B are exhausted.

**Figure 2.**
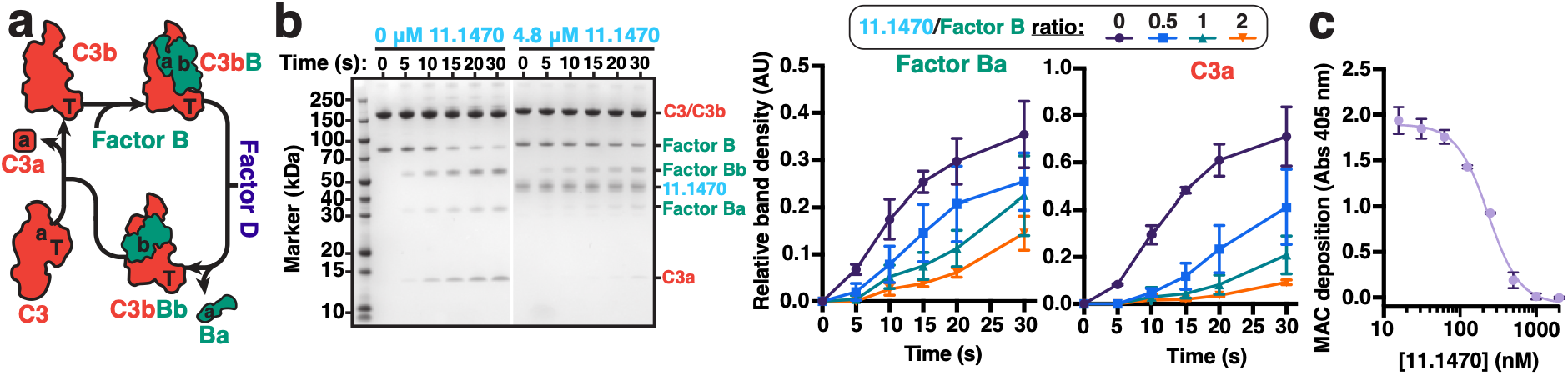
Tb927.11.1470 inhibits C3bBb convertase amplification. (a) Assay for C3bBb convertase amplification. A small amount of C3b is supplied to start the reaction. This binds to Factor B to form C3bB. This pool of C3bB is then cleaved by Factor D, forming C3bBb convertases, which then cleave C3 into C3b. This increases the pool of available C3b molecules to form more C3bBb convertases, and so C3 convertase activity increases over time. (b) On the left, SDS-PAGE showing inhibition of C3bBb convertase amplification by Tb927.11.1470. In the absence of Tb927.11.1470 band density for C3a and Factor Ba increases over time, showing that C3bBb convertase form and cleave C3. Addition of 4.8 μM Tb927.11.1470 (2:1 molar ratio to Factor B) decreases the amount of C3a and Factor Ba observed. On the right, densitometry of SDS-PAGE bands shows that increasing molar ratios of Tb927.11.1470 to Factor B reduces the amount of C3a and Factor Ba formed in a concentration dependent manner. SDS-PAGE is representative of one experimental repeat, with graphs presenting averages from three experimental repeats, n = 3. Tb927.11.1470 concentrations are significantly different, as determined by Two-way ANOVA (For C3a, P = <0.001; for Factor Ba, P = 0.008). (c) Wiesman^TM^ 96-well plate assay for alternative pathway activity, showing inhibition of membrane attack complex (MAC) deposition by Tb927.11.1470. Technical replicates, n = 2. Standard error of the mean is plotted for all graphs.

We repeated this in the presence of Tb927.11.1470 (Figure 2b). The receptor significantly reduced the rates of formation of both C3a and Factor Ba, showing that it inhibits C3bBb amplification (Figure 2b, Supplementary Figure 5a). We repeated this at different ratios of receptor to factor B and, by calculating the area under the curve and fitting a one-phase exponential decay model, we could qualitatively analyse the effect of the receptor (Figure 2b, Supplementary Figure 5b). When equimolar to factor B, Tb927.11.1470 resulted in 70% and 89% inhibition of Factor Ba and C3a production respectively, showing that Tb927.11.1470 is an effective inhibitor of C3bBb amplification.

It is not clear whether the complement cascade progresses to establishment of the membrane attack complex on the African trypanosome surface^25^. Nevertheless, as an assay for C3bBb convertase function, we tested whether Tb927.11.1470 could block progression of the complement cascade to the terminal pathway. We used a Wieslab^TM^ assay, where human serum is added to plates coated with lipopolysaccharide. This activates the alternative pathway, leading to assembly of the membrane attack complex, (MAC) providing a read out of complement pathway progression. In this assay, Tb927.11.1470 inhibited MAC assembly (Figure 2c), with an EC50 of 234 nM, which, assuming a Factor B concentration in serum of 0.5 mg/mL, is equivalent to a molar ratio of 1:1.4 Tb927.11.1470:Factor B. Therefore, two distinct assays indicate that Tb927.11.1470 inhibits amplification of the complement cascade by inhibiting function of the C3bBb convertase.

### Tb927.11.1470 does not prevent Factor B from binding to C3b

While the convertase assay demonstrated that Tb927.11.1470 inhibits amplification of the C3bBb convertase, it did not reveal which step in this cascade was impeded. Given that Tb927.11.1470 binds free Factor B (Figure 1d), we first assessed whether it could prevent Factor B from binding to C3b. We labelled the thioester forming cysteine on C3b with biotin to immobilise on an SPR chip with an orientation which matches C3b conjugation to a pathogen surface. We first studied binding of Factor B to C3b, in the presence of Ni^2+^ to stabilise complex formation^26^, obtaining an affinity of 11 nM (Figure 3a, Supplementary Figure 6, Supplementary Table 3). Tb927.11.1470 binds free Factor B with an approximately 355-fold lower affinity (Figure 1d), suggesting that the receptor will not prevent Factor B from binding to C3b.

**Figure 3.**
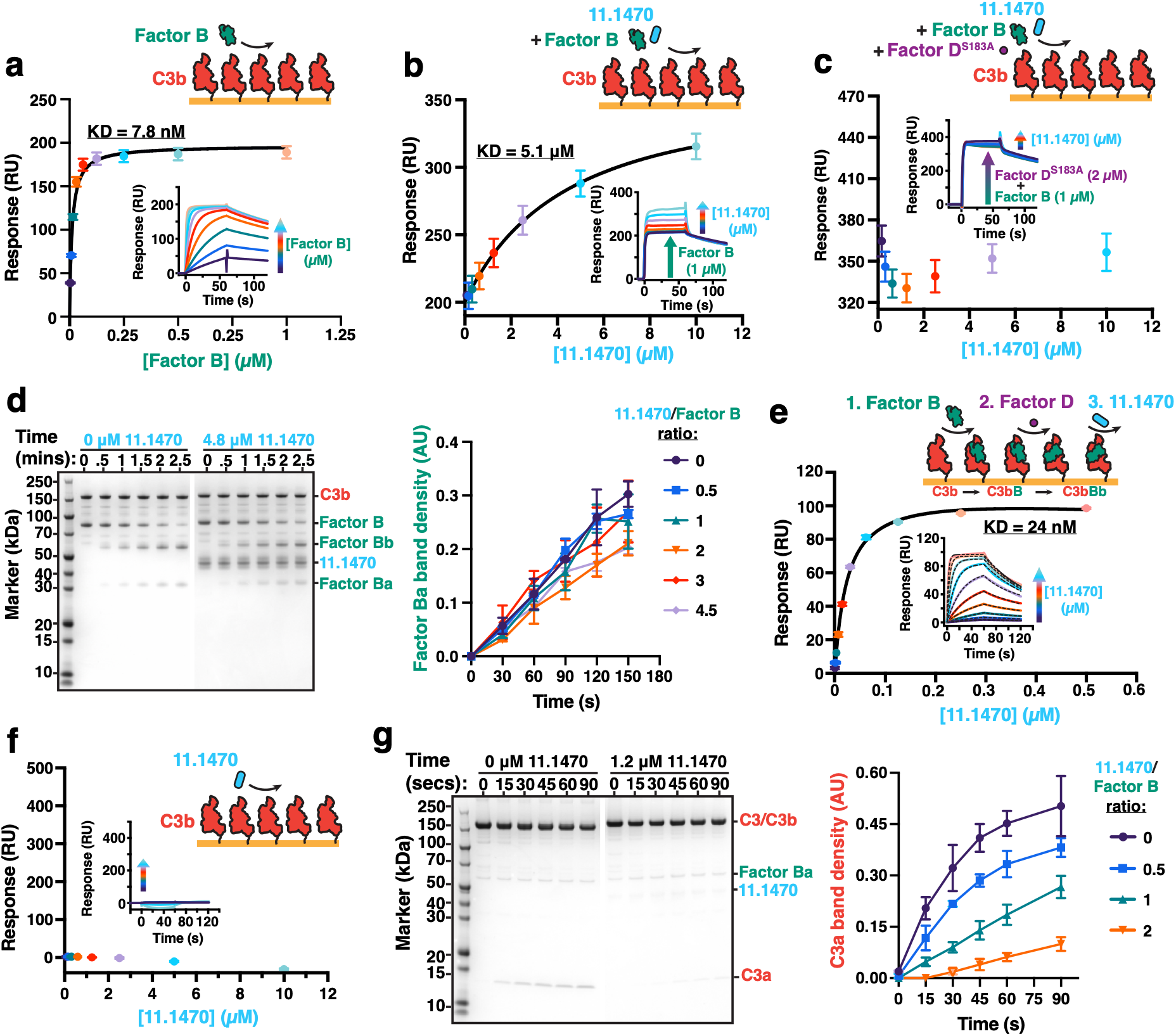
Tb927.11.1470 binds and inhibits the C3bBb C3 convertase. (a) SPR data showing responses from injection of Factor B (two-fold serial dilutions from a concentration of 1 μM) over a streptavidin-coupled flow cell coated with C3b biotinylated on Cys1010. Data is representative of three experiment repeats (n=3). (b) SPR data showing responses from co-injection of Factor B (1 μM fixed concentration) and Tb927.11.1470 (two-fold serial dilutions from a concentration of 10 μM) over a streptavidin-coupled flow cell coated with C3b biotinylated on Cys1010. Data is representative of three experiment repeats (n=3). (c) SPR data showing responses from co-injection of Factor B (1 μM fixed concentration), Factor D^S183A^ (2 μM fixed concentration), and Tb927.11.1470 (two-fold serial dilutions from a concentration of 10 μM) over a streptavidin-coupled flow cell coated with C3b biotinylated on Cys1010. Data is representative of three experiment repeats (n=3). (d) On the left, SDS-PAGE showing that cleavage of C3bB to C3bBb is unaffected by Tb927.11.1470, as shown by similar band density for Factor Ba and Factor Bb in the presence or absence of 4.8 μM Tb927.11.1470 (2:1 ratio of Tb927.11.1470 to Factor B). On the right, densitometry of SDS-PAGE bands shows that increasing molar ratios of Tb927.11.1470 to Factor B does not affect Factor Ba band density. SDS-PAGE is representative of one experimental repeat, with graphs presenting averages from three experimental repeats, n = 3. No Tb927.11.1470 concentrations were significantly different from each other. (e) SPR data showing responses from injection of Tb927.11.1470 (two-fold serial dilutions from a concentration of 0.5 μM) over a flow cell coated with C3bBb. C3bBb is generated from C3b biotinylated on Cys1010 coupled to streptavidin with 1 μM Factor B first injected for 60 seconds, followed by 0.1 μM Factor D for 60 seconds which cleaves Factor B. (f) SPR data showing responses from injection of Tb927.11.1470 (two-fold serial dilutions from a concentration of 10 μM) over a streptavidin-coupled flow cell coated with C3b biotinylated on Cys1010. Data is representative of three experiment repeats (n=3). (g) On the left, SDS-PAGE showing that C3bBb-mediated cleavage of C3 is inhibited by Tb927.11.1470, as shown by a decrease in band density for C3a in the presence of 1.2 μM Tb927.11.1470 (2:1 ratio of Tb927.11.1470 to Factor B). On the right, densitometry of SDS-PAGE bands shows with increasing molar ratios of Tb927.11.1470 to Factor B. SDS-PAGE is representative of one experimental repeat, with graphs presenting averages from three experimental repeats, n = 3. Standard error of the mean is plotted for all graphs. Tb927.11.1470 concentrations are significantly different, as determined by Two-way ANOVA (P = <0.001).

Before Factor B is cleaved, it binds to C3b to form the C3bB pro-convertase. We next determined whether Tb927.11.1470 binds C3bB by injecting a fixed concentration of 1 μM Factor B together with increasing amounts of Tb927.11.1470 over immobilised C3b (Figure 3b, Supplementary Table 4). Increasing the amount of Tb927.11.1470 increased the response over that obtained for Factor B alone, indicating that the receptor can bind C3bB. We calculated an affinity of 4.5 μM for C3bB binding by the receptor, similar to the affinity of the receptor for Factor B alone, indicating that Tb927.11.1470 binds similarly to C3bB and free Factor B.

We next determined whether Tb927.11.1470 prevents binding of Factor D to the C3bB complex using the catalytically inactive Factor D^S183A^ mutant^27^ (Figure 3c, Supplementary Figure 6c). We injected fixed concentrations of 1 μM and Factor B and 2 μM Factor D^S183A^, together with increasing concentrations of Tb927.11.1470. The response was unaffected by the presence of Tb927.11.1470, suggesting that Tb927.11.1470 cannot bind to the C3bB-Factor D^S183A^ complex and that Factor D preferentially binds to C3bB instead of receptor. We therefore predicted that Factor D-mediated turnover of C3bB to C3bBb would occur in the presence of Tb927.11.1470. To test this, we mixed 2.4 μM C3b with 2.4 μM Factor B and 10 nM Factor D and monitored the relative abundance of Factor Ba via SDS-PAGE (Figure 3d, Supplementary Figure 7). Tb927.11.1470 provided at 4.5-fold molar excess over Factor B had no effect on Factor Ba production. Therefore Tb927.11.1470 does not prevent Factor B from binding to C3b and does not prevent its cleavage by Factor D to form the C3bBb convertase.

### Tb927.11.1470 is a C3bBb receptor that inhibits C3bBb activity

Since Tb927.11.1470 did not inhibit C3bBb convertase formation, we hypothesised that it may bind the C3bBb convertase and prevent its C3 turnover activity. We first measured the binding of Tb927.11.1470 to C3bBb using SPR. We bound Factor B to C3b before adding Factor D. This resulted in a drop in response consistent with dissociation of cleaved Factor Ba. We then injected increasing concentrations of Tb927.11.1470, subtracting the response in the absence of receptor to account for Factor Bb dissociation (Figure 3e, Supplementary Figure 8). This revealed that Tb927.11.1470 has a much slower dissociation rate from C3bBb than that observed for its binding to Factor B or C3bB. Indeed, its affinity for C3bBb is 24 nM, which is approximately 250-fold stronger than its affinity for Factor B. We also flowed Tb927.11.1470 over C3b alone and observed no binding (Figure 3f). Therefore, while we initially identified Tb927.11.1470 as a Factor B receptor, it does not block the function of Factor B alone and is, instead, a C3bBb convertase receptor.

We next determined whether binding of Tb927.11.1470 to the C3bBb convertase can inhibit its ability to cleave C3 into C3a and C3b. We reconstituted C3bBb in the presence of increasing amounts of receptor, then added C3 and monitored the relative abundance of C3a as a measure of C3bBb activity (Figure 3g, Supplementary Figure 9). Addition of Tb927.11.1470 potently inhibited C3a production, demonstrating that Tb927.11.1470 directly inhibits C3bBb convertase cleavage of C3.

Therefore Tb927.11.1470 is an inhibitor of the C3bBb convertase, which we name the *T. brucei* C3bBb receptor (C3bBbR).

### C3bBbR stabilises C3bBb in a catalytically inactive form

To determine how C3bBbR inhibits the C3bBb convertase, we determined a cryo-EM structure of the C3bBb convertase in the presence of C3bBbR at 3 Å resolution (Supplementary Figure 10, Supplementary Table 7). This dataset also contained some uncleaved C3bB, allowing us to determine a structure of the open conformation of C3bB at 2.7 Å resolution, compared with 4 Å resolution for the only crystal structure of C3bB^27^ (Supplementary Figure 11). C3bBbR adopts the expected three-helical bundle architecture. It forms interactions with both components of C3bBb, making a 1200 Å^2^ interface with the serine protease domain of Factor Bb, and two smaller 522 and 200 Å^2^ interfaces with the MG1 and TED domains of C3b (Figure 4a). As no interaction was detected for the receptor with C3b at 10μM receptor concentration, these smaller interfaces are insufficient to allow it to bind C3b at physiological concentrations (Figure 3g).

**Figure 4.**
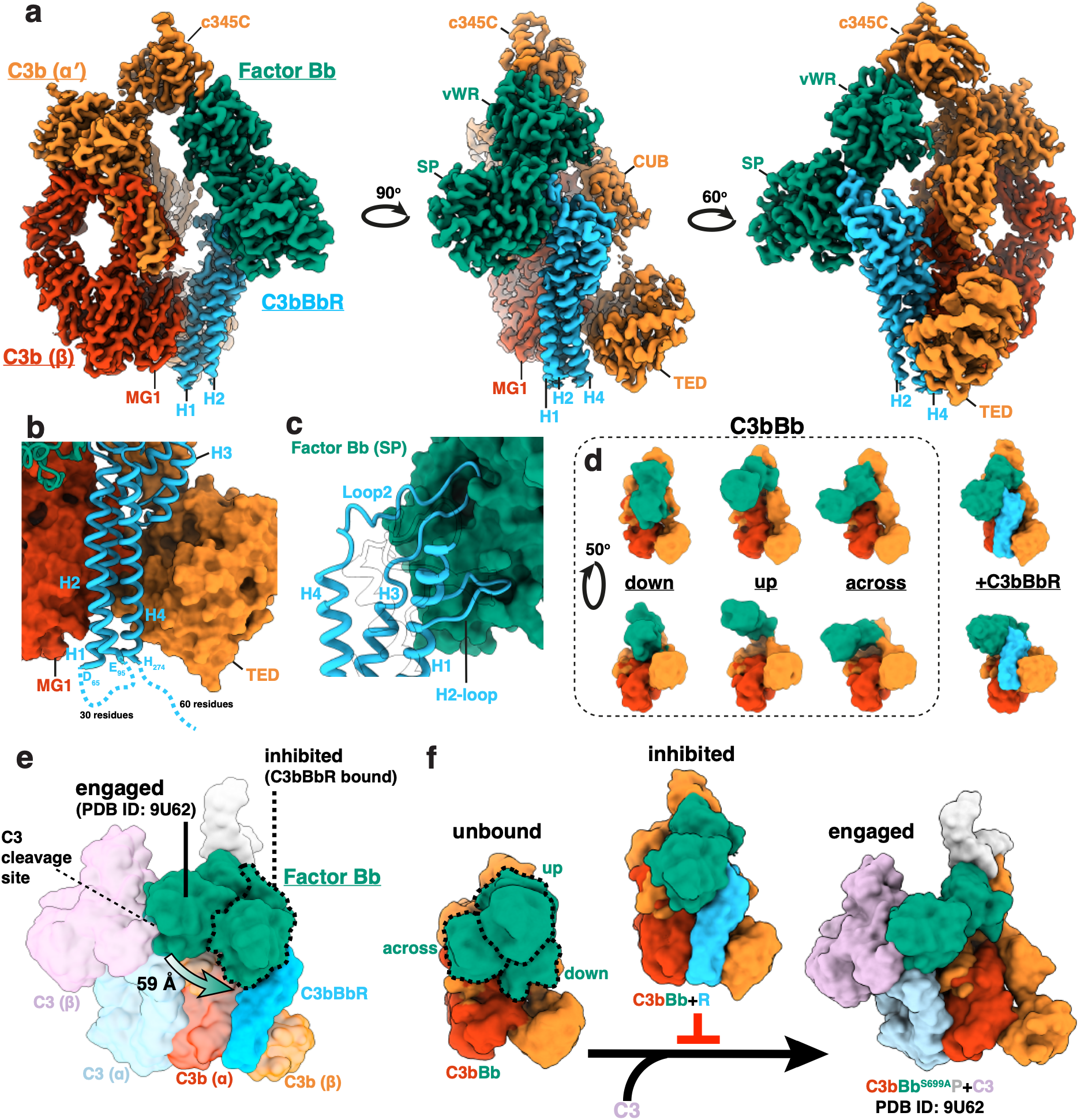
Tb927.11.1470 locks C3bBb in a catalytically inactive conformation. (a) Three views of a 3 Å cryo-EM map of Tb927.11.1470 bound to C3bBb. Tb927.11.1470 (blue), binds the MG1 domain of the C3b β-chain (red), the TED domain of the C3b α-chain (orange) and the serine protease domain (SP) of Factor Bb (green). The helices of Tb927.11.1470 are labelled H1-4. The cryo-EM is a composite map of local alignments focussed on the MG1-8 ring of C3b, and Factor Bb with Tb927.11.1470. (b) Close up of the C3b binding site of Tb927.11.1470, with C3b displayed as a surface, and Tb927.11.1470 as secondary structure ribbon. Blue dotted lines indicate the unresolved regions of the H1, H2, and H4 helices, and the C-terminal tail connecting the 3 helical bundle domain to the transmembrane helix. (c) Close up of the Factor Bb binding site, with Factor Bb displayed as a green surface and Tb927.11.1470 as a blue secondary structure ribbon. The N-terminus, which threads through Loop2, and H2 with connecting loops are displayed transparently with silhouettes. (d) Cryo-EM maps of C3bBb in the presence of magnesium, low-pass filtered to 10 Å. Density for the C3b β-chain is displayed in red, the αʹ-chain in orange, and Factor Bb in green. The top panels present an equivalent view to the middle panel in (a). The right-hand panel shows a surface representation of the C3bBb-Tb927.11.1470 atomic model, displayed at 10 Å. (e) 10 Å resolution surface representation of the structure^28^ of C3 bound to C3bBb^S699A^, superimposed on the structure of C3bBb bound to Tb927.11.1470. Factor Bb from the Tb927.11.1470 bound structure is highlighted with a dotted line. (f) Model of a C3bBb inhibition by Tb927.11.1470. The left shows an overlay of the three conformations in (d), showing three conformations of factor B (highlighted with dashed lines). The central panel shows the receptor bound conformation of C3bBb, showing factor B in a downwards conformation, while the left panel shows C3bBb with factor B in an across conformation to engage a C3 substrate. By preventing the adoption of the across conformation of factor B, the receptor blocks C3bBb function.

The bottom half of the three helical bundle sits in the opening between the Macroglobulin 1 (MG1) and thioester (TED) domains of C3b, with Helix1 of C3bBbR packed against the central beta-sheet of MG1, and Helix4 forming a salt bridge with C3d (Figure 4b, Supplementary Figure 12a and b). Two loops at the top of C3bBbR mediate the interaction with Factor B: the 30 amino acid Loop2 which connects helices 3 and 4, and a short loop that emerges from Helix2 (H2-loop) (Figure 4c, Supplementary Figure 12c). The two loops bind distinct grooves in Factor B, forming a clamp around the serine protease domain. Thus, by binding to both C3b and Factor Bb, C3bBbR locks C3bBb in a defined conformation.

To understand the impact of the C3bBbR induced conformation on C3bBb function, we used cryo-EM to investigate the structure of C3bBb in the presence of magnesium, solving a 2.8 Å consensus structure (Supplementary Figure 13). The density for Factor Bb is blurred because it is bound only to the small C345C domain of C3b, which is flexible relative to the MG1-8 ring of C3b. We therefore applied 3D classification and resolved several states to represent different snapshots of the continuum of Factor Bb locations. We chose three representative conformations that we categorise as ‘across’, ‘down’, and ‘up’ (Figure 4d). In the down state, the serine protease domain of Factor Bb is located between the top of the TED and MG1 domains of C3b, overlapping with the C3bBbR binding site, whilst in the up-state Factor Bb is located away from the body of C3b. Whilst no classes aligned exactly with the Factor Bb conformation that we observed in the C3bBbR structure, the across state of C3bBb is most similar. Overall, this data shows that C3bBbR does not stabilise a dominant conformation of C3bBb. However the range of Factor Bb motion in unbound C3bBb overlaps significantly with the conformation observed in C3bBbR, which may contribute to efficient binding of the receptor.

A recent cryo-EM structure used catalytically inactive Factor B to reveal the conformation of C3bBb bound to properdin, a positive regulator that stabilises C3bBb, when bound to its C3 substrate^28^. This showed Factor Bb when engaged with the scissile 746-747 bond in a cleavage ready state. In this structure, the MG1-8 rings of C3b and C3 dimerise, enabling Factor Bb to engage with the scissile bond on C3. Superimposing the structure of C3bBbR bound to C3bBb reveals that the receptor holds Factor Bb 59 Å away from the C3 engaged conformation (Figure 4e). This differs from the previously characterised C3bBb regulator SCIN, which binds to the C3b MG6-7 domains at the top of the MG-ring, and the MG7-8 domains on a second C3b molecule to form an inactive dimer^16^. C3bBbR therefore functions by stabilising a catalytically incompetent C3bBb conformation which is unable to swing across to the location required to engage a C3 substrate.

## Discussion

Here we discover C3bBbR, a novel complement regulating protein on the surface of the African trypanosome *Trypanosoma brucei*. Originally, we identified this protein to bind with micromolar affinity to Factor B, however our subsequent characterisation showed C3bBbR to be a high affinity binder of C3bBb, the C3 convertase that amplifies the complement cascade. We also analysed the cryo-EM structure of unbound C3bBb, showing Factor Bb to adopt a continuous but defined range of conformations. Upon binding of C3bBb to its C3 substrate, Factor Bb must reach across the MG ring of C3b to engage with the scissile bond of C3. Notably, this conformation is outside the range of conformations we identified in unbound C3bBb, so is not a major conformation in the absence of C3 (Figure 4f). Upon binding of C3bBbR, Factor Bb is locked in a conformation that prevents it from reaching the scissile bond of C3, directly inhibiting its convertase function.

Whilst targeting the C3bBb convertase does not prevent initiation of the complement system, it prevents amplification of C3b deposition. C3bBb inhibition therefore has the potential to prevent accumulation of C3b-based opsonins on the pathogen surface to levels where complement effectors would be operational. Other pathogens have also evolved to target the C3bBb convertase, most notably *Staphylococcus aureus,* which uses SCIN to inhibit C3 convertases^16^ and leads to significant reduction of phagocytosis of bacteria^29^. SCIN is able to bind to C3b molecules simultaneously, resulting in stable dimers that are catalytically incompetent. The mosquito *Anopheles albimanus* has evolved a structurally similar protein to SCIN, albicin, that also binds to the top of the MG-ring and results in stable, inactive C3bBb and C3b dimers^30^. These proteins also form interactions with the von Willebrand domain of Factor Bb. Unlike C3bBbR, SCIN and albicin also bind free C3b, driven by a low affinity binding site on the serine protease domain of Factor B that combines with binding to the bottom of the MG-ring on C3b to form a high affinity interaction. Binding to the bottom of the MG-ring, which is parallel to the thioester bond in the TED, is presumably compatible with the surface tethered nature of C3bBbR. Recently, CirpA1 from the hard tick *Rhipicephalus pulchellus* was discovered which prevents the C3bBb stabilising protein properdin from binding to C3bBb^31^, therefore indirectly reducing C3bBb-based amplification. In contrast, C3bBbR is the first pathogen protein known to selectively bind to C3bBb convertase and to function by bridging domains of C3b and Bb to stabilise a catalytically incompetent Factor Bb conformation.

As an extracellular parasite, the cell surface of African trypanosomes is under extraordinary pressure from the immune system. Much like the bacterial pathogen *S. aureus^16,32–34^*, African trypanosomes simultaneously target multiple complement components. With three complement receptors now discovered^9,22,23^, we have begun to understand how this pathogen surface has evolved to counteract the threat of complement. These receptors share a structurally similar three-helical bundle architecture but mediate distinct complement evasion strategies. The Factor H receptor, which was recently found to be expressed in both bloodstream form trypanosomes^35^ and the procyclic insect form^9^, recruits host Factor H, which negatively regulates complement by accelerating C3bBb convertase decay amongst other functions. This is a very common strategy among pathogens^36^, and the Factor H receptor may work in tandem with C3bBbR to inhibit C3bBb amplification. In contrast to C3bBbR, ISG65 binds directly to C3b, and whilst a variety of mechanisms for this protein have been suggested, it is not clear how it functions in the context of the parasite ^22,23,37,38^. This shows that African trypanosomes have evolved receptors which target different aspects of the complement system, potentially allowing them to act synergistically.

The discovery of C3bBbR reveals a novel and potent inhibitor of the C3bBb convertase on the surface of African trypanosomes. C3bBbR stabilises C3bBb in a catalytically inactive conformation, and its binding specificity to C3bBb may be beneficial in design of therapeutically useful complement inhibitors. It remains unclear how this receptor synergises with other *T. brucei* complement binding proteins and how they come together to mediate complement resistance. It shows that African trypanosomes have evolved a broad network of complement inhibitors which target different stages of the complement cascade and which has yet to be fully uncovered.

## Acknowledgements

This work was funded by Wellcome Discovery Awards (217138/Z/19/Z and 220797/Z/20/Z). We thank Rishi Matadeen, Courtney Mycroft-West, and Edward Lowe at the COSMIC facility (University of Oxford) for support with cryo-EM data collection and data processing and David Stauton for support with biophysics. We thank Hannah Ivison for laboratory management.

## Author contributions

A.D.C. designed, conducted and analysed experiments. N.M., H.W. and M.C. conducted bioinformatics, provided expertise in trypanosome receptor selection and provided cloned receptors. M.C. and M.K.H. contributed to experimental design and provided expertise and funding. A.D.C and M.K.H. prepared the manuscript and all authors contributed and commented.

## Data and materials availability

Cryo-EM maps and associated co-ordinates for the C3bBbR:C3bBb structure are available as follows. Protein Data Bank (PDB) coordinates (33DX); Electron Microscopy Data Bank (EMDB) accession codes: composite map (EMDB-59383), consensus map (EMDB-59379), local alignment on Factor Bb-C3bBbR (EMDB-59383), local alignment on MG-ring of C3b (EMDB-59382). The C3bBb only reconstructions are available on the EMDB as: ‘up’ conformation (EMDB-59387), ‘down’ conformation (EMDB-59388), and ‘across’ conformation (59392). The C3bB structure is available as: PDB coordinates (33EG), EMDB (EMDB-59391). All other raw data is available in the Source Data file and materials are available from the authors on request.

## Materials and methods

### Selection of a panel of putative T. brucei cell surface receptors

A panel of putative receptors was selected using sequence data available at the time (2019). First, we included receptors whose cell surface localisation was predicted by either the presence of a likely GPI-anchor signal using NetGPI 1.1^39^ or a single transmembrane helix using TMHMM^40^. Second, we only included proteins with mature extracellular domain >20kDa as this was considered to be the size necessary to form a folded domain which binds a protein ligand in the context of the VSG coat. Third, we only included proteins for which transcriptome data indicated expression in bloodstream form trypanosomes. The highly abundant cell surface proteins characteristic of each life cycle stage; VSG^17^, procyclins^41^, BARP^42^ and MISP^43^ were excluded from the final list.

The screen was then supplemented by searching for invariant surface glycoproteins (ISGs) known to be cell surface localised^22,44^, by using Blast to iteratively search the proteome with previously identified ISGs and then newly identified homologues until no further candidates were identified. A similar approach was taken with proteins with a three helical bundle architecture as this is a common motif present in trypanosome receptors^45^.

The transferrin receptor is structurally similar to VSGs^21^ and a small number of proteins adopting the VSG fold were included in the list if they met the following criteria. First, that the sequence was present and conserved in distantly related isolates with available genome sequences at www.TriTrypBD.org: *T. b. brucei* TREU927 and *T. b. brucei* Lister427 and *T. b. gambiense* Daloa972. Second, if the gene was located near the end of a polycistronic transcription unit likely to be transcribed by RNA polymerase II and so located in a chromosome core^46^. Third, if expression was detected in a transcriptome analysis in blood stream form trypanosomes.

### Expression and purification of a T. brucei surface protein library

The ectodomains of all identified putative surface proteins were cloned into a pCI plasmid with an N-terminal CD33 secretion signal and C-terminal GSGSGSSNSGSRASG linker, AviTag^TM^, and His_10_-tag. Ectodomain amino acid boundaries are denoted in Supplementary Table 1. To produce biotinylated protein, surface proteins were co-expressed in HEK293F cells with BirA (Addgene catalog #64395). 2.5×10^6^/mL of HEK293F cells were pelleted and resuspended in an equivalent volume of fresh Freestyle F17 media 100 μM D-biotin. 3 ug of *T. brucei* surface DNA, and 0.3 ug of BirA DNA per mL of cells were added, followed by incubation at 37°C whilst rotating for 5 minutes. 9 ug of PEI per mL of cells was then added. After incubation for 20 hours, the cell culture volume was then doubled and 3.8 mM valproic acid added. After 5 days, cell supernatants were harvested and applied to a 1 mL HisTrap^TM^ Excel column (Cytiva) using an ÄKTA Pure 25 (Cytiva). The column was washed with IMAC wash buffer (20 mM Tris pH8, 500 mM NaCl, 10 mM imidazole), and bound protein eluted with IMAC elution buffer (IMAC wash with 300 mM imidazole). The elution was concentrated using Amicon Ultra centrifugal filters, resuspended in HBS (HEPES buffered saline: 20 mM HEPES pH 7.4, 150 mM NaCl), and concentrated again.

Proteins with little or no expression were cloned into an pExpesS2 vector containing an N-terminal BiP secretion signal and C-terminal His_6_-tag, and stable *Drosophila* S2 cell lines generated. S2 cells were transfected as static cultures at 25°C using ExpreS^2^ Insect-TR transfection reagent (ExpreS^2^ion Biotechnologies) and cell lines were selected with Zeocin (ThermoFisher). For proteins with low expression, HEK293F and S2 culture volumes were increased accordingly (Supplementary Table 1).

### Source and Purification of Complement components

Total human IgM, C-reactive protein, and serum amyloid P were purchased from Sigma (product codes I8260, 236603, 555145 respectively). Complement C1, C4, C4b, C5, C5b6, C6, C7 C8, C9, Factor I, and C4 binding protein were purchased from Complement Technologies (product codes A098, A105, A108, A120, A122, A123, A124, A125, A126, A138, A109 respectively).

To purify Complement C3, anonymous donor post-clot human serum was obtained from the NHS Blood and Transplant non-clinical issue supply. Serum was centrifuged at 12,000 g for 15 mins at 4°C and then passed through a 0.45 μm filter. 2% (w/v) polyethylene glycol 4000 (PEG4000) was added and the mixture was gently agitated for 30 mins at 4°C. The serum was centrifuged for 30 mins at 10,000 g, the supernatant was retained and 7% PEG4000 was added as above, followed by centrifugation. The pellet was washed twice with PBS, dissolved in Q wash (20 mM Tris pH 8.8, 80 mM NaCl, 0.5 mM EDTA) and loaded onto a 1 mL Resource Q column (Cytiva). The column was washed with Q wash, then eluted with Q elution (Q wash + 1 M NaCl) over a 0-30% gradient. Fractions containing C3 were pooled and exchanged into S wash (20 mM Acetate pH 5, 80 mM NaCl, 0.5 mM EDTA) using a Centripure P100 column (emp BIOTECH), then loaded onto a 1 mL Resource S column (Cytiva). The column was washed with S wash, then eluted with S elution buffer (S wash buffer + 1 M NaCl), over a 30% gradient. Fractions containing C3 were pooled, concentrated, and further purified on a Superdex 200 300/10 GL (Cytiva).

To generate C3b from C3, 1:100 w/w trypsin (Roche) was added for 2 mins at 37°C, then 2:100 w/w soybean trypsin inhibitor (Merck) immediately added. For biotinylation of C3b, 100 mM HEPES pH 7.0 was added after addition of soybean trypsin inhibitor, followed by a 10-fold molar excess of maleimide-PEG2-biotin (ThermoFisher). The reaction was incubated on ice for 6 hr. C3b or biotin-C3b was then purified on a Superdex 100 300/10.

Complement C2, Factor D, Factor B were cloned into the pHL-sec vector with C-terminal GSG linker and C-tag, whilst properdin was cloned into pHLsec with C-terminal his_6_-tag. C2, Factor D, and properdin were expressed in HEK293F cells as above, whilst Factor B was expressed in Expi293F GnTi^-^ cells (ThermoFisher) as per the manufacturer’s instructions. C2, Factor D, and Factor B were purified using CaptureSelect^TM^ C-tagXL resin (ThermoFisher) and eluted with 20 mM Tris pH 8, 2 M MgCl_2_, whist properdin was purified using Nickel Excel resin as above. The catalytically inactive Factor D^S183A^ mutant was expressed and purified as for Factor D. Factor Ba and Factor Bb fragments were expressed and purified as for Factor B. Total IgG was purified from human serum using protein A/G resin (Cytiva), and eluted using 100 mM glycine pH 2.8 into tubes containing 1 M Tris pH 8. Eluate was concentrated and further purified on a Superdex 200 300/10 GL.

C3d^C1010A^ and C4d^C1010A^ were expressed in BL21 *E. coli* (NEB) in a pET51b vector with a N-terminal His_6_-tag and TEV cleavage site. *E. coli* were grown in 2xYT broth containing ampicillin at 37°C, shaking at 200 rpm to an optical density at 600 nm of 0.6 and expression was induced with 1 mM Isopropyl β-D-1-thiogalactopyranoside. Cultures were incubated overnight at 18°C then lysed using a CF2 cell disruptor (Constant Systems). The lysate was clarified by centrifugation at 40,000 g, then C3d/C4d purified by nickel affinity chromatography. Nickel Eluate then concentrated and C3d/C4d further purified on a Superdex 75 10/300.

### Screen of trypanosome and complement libraries

Putative *T. brucei* receptors were tetramerised by addition of Streptavidin-HRP (21124, ThermoFisher) at a 4:1 molar ratio. Tetramerised receptors were stored in 96-well plates at −70°C after snap-freezing in liquid nitrogen. Prior to use, samples were defrosted and passed through 0.2 μm filter plates (Agilent).

Complement components were non-specifically immobilised on MaxiSorb^TM^ 96-well plates (Nunc) at 4 μg/mL in 100 mM NaCO_3_ pH 9.3 for an hour. Wells were washed three times with HBS + 2 mM MgCl_2_ + 0.05% tween-20 (HBST-Mg) using a plate washer, then 300 μL Blocker^TM^ Casein (37528, ThermoFisher) was added and incubated for an hour. The plate was then washed three times using HBST-Mg without Tween-20 (HBS-Mg). 7 ug/mL tetrameric prey diluted in Blocker^TM^ Casein was then added and incubated for an hour. For properdin, it was necessary to add 100 μM biotin to prevent non-specific binding. The plate was then washed 3 times with HBST-Mg, once with HBS-Mg and was then developed by adding 50 μg/mL TMB in 100 mM Citrate-phosphate pH 5.3. After 10 mins, 50 μL 1 M sulphuric acid was added to block the reaction, and absorbance values read at 450 nm using a ClarioStar (BMG LABTECH). Background signal was subtracted using wells with only streptavidin-HRP added. For each complement component, C3 and C4d were also immobilised and ISG65 added to both to provide a positive and negative control. Streptavidin-HRP with no receptor was also added as a negative control.

### Solution C3bBb convertase assays

To assess the effect of Tb927.11.1470 on C3bBb amplification, 6.5 μM C3, 2.4 μM Factor B, 70 nM C3b, and 70 nM Factor D were combined in a 20 μL reaction with 0, 1.2, 2.4, or 4.8 μM Tb927.11.1470 in HBS-Mg. Firstly, C3, Factor B, C3b, and Tb927.11.1470 were combined in 10.5 μL at a 2X concentration, and a 0.5 μL sample removed at time point 0 seconds. 10 μL of Factor D at 2-fold concentration was then added, and 1 μL samples collected at 5, 10, 15, 30, and 45 seconds, with the reaction quenched by adding SDS-PAGE loading buffer (ThermoFisher, NP0007)

To assess the effect of Tb927.11.1470 on Factor B cleavage by Factor D, 2.4 μM C3b and 2.4 μM Factor B, D were combined in a 20 μL reaction with 0, 1.2, 2.4, or 4.8 μM Tb927.11.1470 in HBS-Ni. 1 μL sample was removed at time point 0 seconds. 7 nM Factor D was then added, and time points were taken at 30, 60, 90, 120, and 150 seconds, with the reaction quenched by adding SDS-PAGE loading buffer.

To assess the effect of Tb927.11.1470 on C3bBb cleavage of C3, 0.6 μM C3b, 0.6 μM Factor B, were combined in a 20 μM reaction with 0.3, 0.6, or 1.2 μM Tb927.11.1470. in HBS-Mg. 0.1 μM Factor D was then added, incubated for 20 seconds, then 6.5 μM C3 added and a sample immediately added to SDS0PAGE sample buffer. Further time points were then collected at 15, 30, 45, 60, and 0 seconds, with the reaction quenched by adding SDS-PAGE loading buffer.

Samples were run on NuPAGE^TM^ 4-12% Bis-Tris Midi gels, and stained in Quick Coomassie Stain (Generon) overnight. Gels were washed three times with distilled water, then imaged on an iBright FL1500 for 10 seconds. Densitometry was performed in FIJI^47^. Where no density for a band was observed, a value of zero was used.

### Surface plasmon resonance analysis

To validate binding of Tb927.11.1470 to Factor B-derived molecules, 10 μg/mL Factor B, Factor Ba, or Factor Bb were immobilised on a CM5 chip (Cytiva) in 50 mM acetate pH 4 to 510, 249 or 464 responses. Tb927.11.1470 was buffer exchanged into SPR buffer (20 mM HEPES pH 7.4, 150 mM NaCl, 0.005% tween-20) using Zeba^TM^ spin 0.5 mL 7 MWCO columns (ThermoFisher). A seven step two-fold dilution series was prepared from 40 μM Tb927.11.1470, which was injected with a flow rate of 30 μL/min for 120 seconds. Dissociation was then allowed to occur for 120 seconds before regenerating the chip for 60 seconds with 20 mM acetate pH 4, 500 mM NaCl between each sample injection.

To investigate Tb927.10.3620 binding to C3, Tb927.10.3620 was conjugated to streptavidin-HRP and immobilised as above to 988 RU on a CM5 chip. In addition, 781 RU streptavidin was immobilised, followed by 221 RU of an independent purification of biotinylated Tb927.10.3620, as well as 771 RU streptavidin followed by 206 RU of biotinylated ISG65 as a positive control. A seven step two-fold serial dilution of 2 μM C3, C3b, and C3d was prepared and injected at 30 μL/min for 60 seconds, with dissociation of 60 seconds, with regeneration as above.

For binding of Factor B to C3b, 602 RU streptavidin was immobilised on a CM5 chip. 478 RU of C3b biotinylated on Cys^1010^ was then added. An 8 step 2-fold serial dilution of 1 μM Factor B was then prepared and injected at 30 μL/min for 60 seconds, with dissociation of 60 seconds, then regenerated for 100 seconds with 20 mM acetate pH 4, 500 mM NaCl, 20 mM EDTA.

For binding of Tb927.11.1470 to C3bB and C3bBD^S183A^ a seven step two-fold serial dilution of 10 μM Tb927.11.1470 was prepared, and 1 μM Factor B alone or in combination with 2 μM Factor D^S183A^ was added to all dilutions. These samples were injected over immobilised C3b as for Factor B above. For binding to C3bBb, 1 μM Factor B was first injected, followed by injection of 0.5 μM Factor D, which cleaved C3bB to yield C3bBb on the chip. A nine step two-fold serial of 0.5 μM Tb927.11.1470 was then injected. The regeneration solution was sufficient to remove Tb927.11.1470 but not Factor Bb from C3b. Factor B and Factor D were therefore added before each Tb927.11.1470 injection, to ensure equivalent levels of C3bBb on the chip.

Steady state parameters were estimated using GraphPad Prism 11, and kinetic parameters measured using Biacore Evalution v1.0.

### Preparation of C3bBb-Tb927.11.1470 complexes and imaging with Cryo-EM

C-Flat grids composed of 300 mesh copper with a 1.2/1.3 uM holey carbon film were glow discharged at 15 mA for 1 min with an EM ACE200 glow discharger (Leica). C3b, Factor B, and Tb927.11.1470 were pooled at a 1:1:1.1 ratio in 20 mM HEPES pH 7.4, 150 mM NaCl, 2 mM NiCl_2_ to a final concentration of 2.5 mg/mL. 0.1 nM Factor D was added for 30 seconds, followed by addition of 0.01% fluorinated octyl maltoside (Anatrace), and immediate transfer to grids and plunge freezing with a Vitrobot Mark IV (Thermo Fisher). Grids were imaged on a Titan Krios G2 (Thermo Fisher) operating at 300 kV and images recorded on a K3 detector (Gatan) in counting mode in conjunction with a BioQuantum 20 eV energy filter (Gatan). 48,680 movies we recorded at 105,000x magnification with a pixel size of 0.83 Å/pix and a total does of 40.3 e^-^/Å^2^ over 40 frames.

C3bBb grids were prepared and imaged as above except: C3b, Factor B and Factor D were pooled at a 1:1:0.1 ratio to a final concentration of 2.5 mg/mL in 20 mM HEPES pH 7.4, 150 mM NaCl, and 2 mM MgCl_2_. Proteins were incubated for 20 seconds, followed by addition 0.01% fluorinated octyl maltoside and immediate plunge freezing. 28,057 movies were recorded.

### Image processing of C3bBb-Tb927.11.1470 and C3bB complexes

Image processing was performed in CryoSPARC v4 (Structura Biotechnology Inc.)^48,49^. Movies were pre-processed using patch motion correction and patch contrast transfer function (CTF) estimation. For particle picking, template matching using circular/elliptical blobs between 100-190 Å diameter was performed on 500 micrographs. Three rounds of 2D classification yielded five clear 2D classes which were used in a second round of template matching before another three rounds of 2D classification yielded eight clear classes. Temple matching using these classes was then performed on 5746 micrographs, and a single round of 2D classification using 200 classes was performed and 200,322 particles were selected. Ab initio reconstruction was then performed with the five classes, follow by heterogenous refinement using all resultant classes and particles from *ab initio*. 84,933 particles corresponding to a single class with clear C3b density were then selected and subjected to a second round of Ab initio reconstruction followed by heterogeneous refinement, which yielded a clear class for C3bBb bound to Tb927.11.1470, a clear class corresponding to C3bB, and two further junk classes. 69,679 particles belonging to the C3bBb-Tb927.11.1470 and C3bB classes were then pooled, and a TOPAZ model^50^ was developed on the picks from 95 micrographs using an expected particle number of 70 over 15 epochs. In addition, 2D classification from the pooled particles yielded a final set of 26 classes for template matching.

The 2D templates and the TOPAZ model were then used to pick particles on all micrographs, yielding 3,528,997 and 2,030,324 particles. These were subjected to a single round of 2D classification with 200 classes, followed by two rounds of heterogeneous refinement using the references generated above. Particles from template and TOPAZ picking were merged and duplicates removed, yielding 466,906 particles corresponding to C3bBb-Tb927.11.1470 and 859,496 to C3bB.

For C3bBb-Tb927.11.1470, non-uniform refinement with per-particle scale and CryoSPARC implementation of per-particle defocus estimation^51^ yielded a 3 Å map. The CryoSPARC implementation of reference-based motion correction^52^ subsequently yielded a 2.9 Å map. To improve the occupancy of Factor Bb and Tb927.11.1470, Segger^53^ was used to segment density corresponding to Tb927.11.1470, Factor Bb, and the C345C domain of C3b, which was used to perform focussed 3D classification with five classes, limited to 10 Å resolution. Three classes consisting of 287,671 particles were selected, yielded a 3 Å map after non-uniform refinement. Particles had a bimodal distribution of per-particle scale values, and a subset of 164,123 particles with values between 0.96 and 1.92 were selected, yielding a 2.9 Å map. The serine protease domain of Factor Bb suffered lower local resolution in some parts, owing to limited mobility of Factor Bb, Tb927.11.1470, and the C3b C345C domain relative to the main ring of C3b. Therefore, Segger was used to generate a mask (soft edge of 12 pixels) corresponding to the ring of C3b (domains MG1-8), which was used in a focussed alignment (using a pose/shift gaussian prior during alignment, a standard deviation of prior over rotation of 7°, and a standard deviation of prior over shifts of 3.5 Å) to generate a 2.8 Å map. The mask was then used to subtract the signal for MG1-8, and local alignment using a mask (soft edge of 12 pixels) covering C345C, Factor Bb, and Tb927.11.1470 preformed, generating a 3 Å map. Phenix^54^ was used to generate composite maps from local alignments.

For C3bB, non-uniform refinement with per-particle scale, per-particle defocus, and reference motion correction yielded a 2.6 Å map. Density corresponding to Factor B only was extracted and mask 12-pixel soft edge mask generated for focussed 3D classification (resolution = 4 Å, number of O-EM epochs = 20, number of classes = 10). 142,437 particles with improved occupancy of Factor B were selected, yielding a 2.8 Å map. Map post-processing was performed using DeepEMhancer^55^.

### Model building of C3bBb-Tb927.11.1470 and C3bB

As a starting model for C3bBb-Tb927.11.1470, crystal structures of C3b^11^ (5FO7) and Factor Bb^56^ (1RRK), and an Alphafold3^56^ model of Tb927.11.1470 were rigid body fitted into the map using ChimeraX^57^. To flexibly fit the serine protease domain of Factor B and the C345C domain of C3b into the map, distance restraints were applied to all atoms, and an Isolde^58^ simulation in ChimeraX^59^ was run until the domains settled into density. For C3bB, an Alphafold3 model was generated and used as a starting model. Isolde was then used to rebuild incorrect loops, adjust sidechain and backbone geometry, and add glycans where appropriate. Density corresponding to the CUB and C3d domains of C3b was poor compared to the rest of map, owing to flexibility. Therefore the 2.8 Å crystal structure of C3b^11^ (5FO7) was used to generate reference-based restraints in Isolde for these regions. A combination of the unsharpened map and DeepEMhancer postprocessed map was used with Isolde. After each round of model building in Isolde and Coot^60^, Phenix real-space^61^ refine was performed against the unsharpened map, until no further improvements could be made.

### Image processing of C3bBb

Movies were pre-processed as above. For particle picking, template matching using circular and elliptical blobs between 90 and 190 Å in diameter was first performed on 2,285 micrographs. 2D classification with 200 classes was then performed, and 10 classes used as templates for particle picking of the entire dataset, yielding 4,369,630 initial particles. Three rounds of 2D classification using 200 classes were then performed yielding 954,146 particles. *Ab initio* reconstruction followed by heterogeneous refinement with 5 classes was then performed, yielding two classes with clear C3b density, corresponding to 648,931 particles. After reference-based motion correction, homogeneous refinement with per-particle defocus optimisation and per-particle scale minimisation yielded a 2.7 Å map.

Factor Bb contacts the C345C domain of C3b, which is very mobile relative to the rest of C3b. Therefore C345C and Factor Bb density was very blurred in this reconstruction. To obtain reconstructions with improved Factor Bb density, and to removed unbound C3b particles, 3D classification in RELION v5.0 was performed^62^. Firstly, 3D auto-refine was performed, yielding a 2.8 Å map. To generate a focussed mask for C345C and Factor Bb that encompassed the range of movements, spherical and elliptical masks were generated and summed in ChimeraX. 3D classification with alignment and 40 classes was then run, yielding 226,715 particles with clear Factor Bb density. To generate a more accurate mask, the volumes from these classes were summed, and density corresponding to C345C and Factor B extracted. A second round of 3D classification with 20 classes was then performed yielding 11 classes with clear Factor Bb density. Three representative classes of different conformations were picked and subjected to non-uniform refinement in CryoSPARC. The range of Factor Bb motion was continuous, therefore the resolution of Factor Bb was always significantly lower than for the MG1-8 domains of C3b. All reconstructions were therefore low pass filtered to 10 Å.

### Graphing and molecular visualisation

Molecular graphics and analyses performed with UCSF ChimeraX^57,59^, developed by the Resource for Biocomputing, Visualization, and Informatics at the University of California, San Francisco, with support from National Institutes of Health R01-GM129325 and the Office of Cyber Infrastructure and Computational Biology, National Institute of Allergy and Infectious Diseases. Graphs were plotted and analysed in GraphPad Prism v11.

**Supplementary Data: Bioinformatic screen for putative *T. brucei* cell surface receptors, nd = not determined, TPM = Transcripts Per Kilobase Million.**

## Supplementary Figures

**Supplementary Figure 1.**
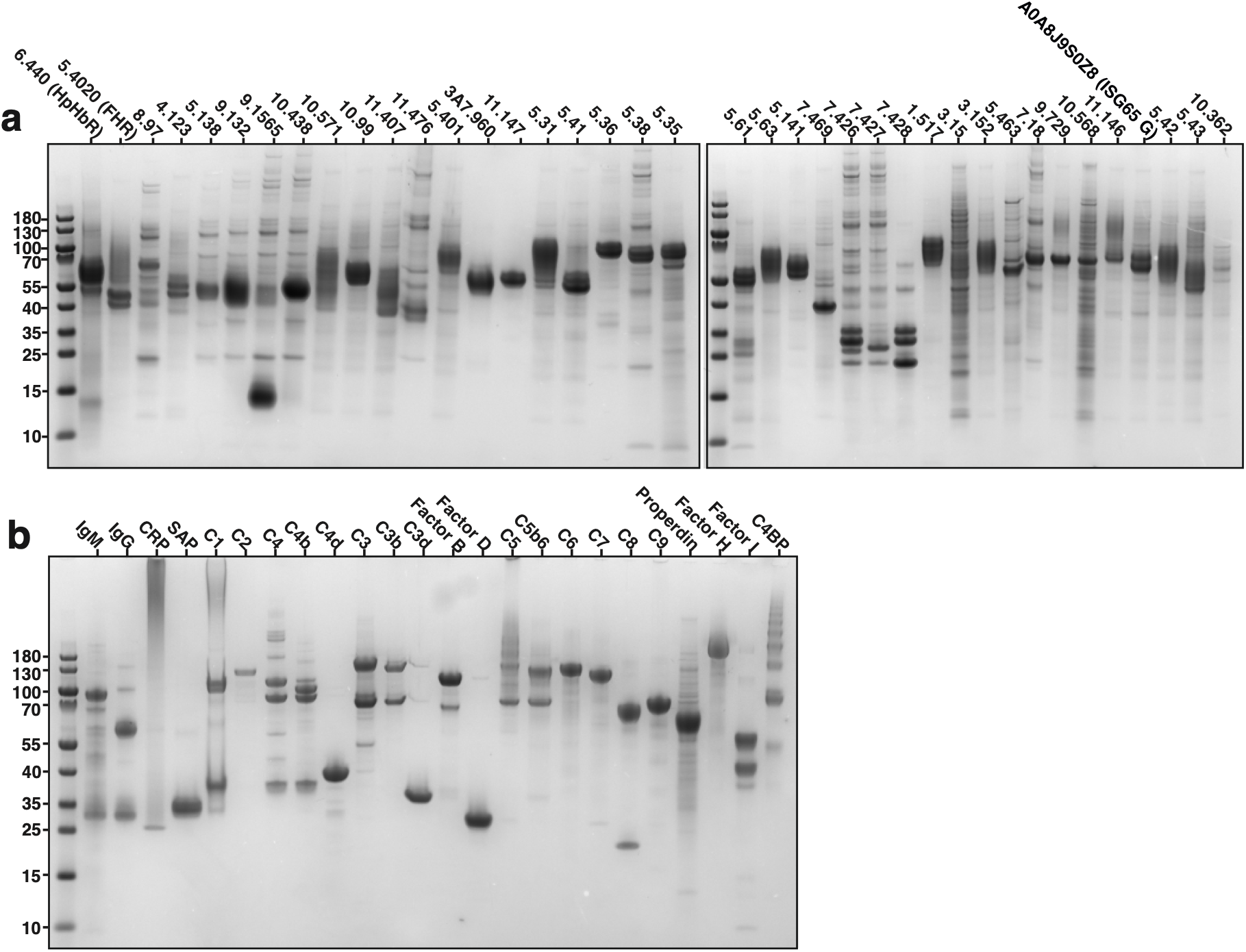
SDS-PAGE gels of purified (a) putative *T. brucei* surface proteins, (b) complement proteins.

**Supplementary Figure 2.**
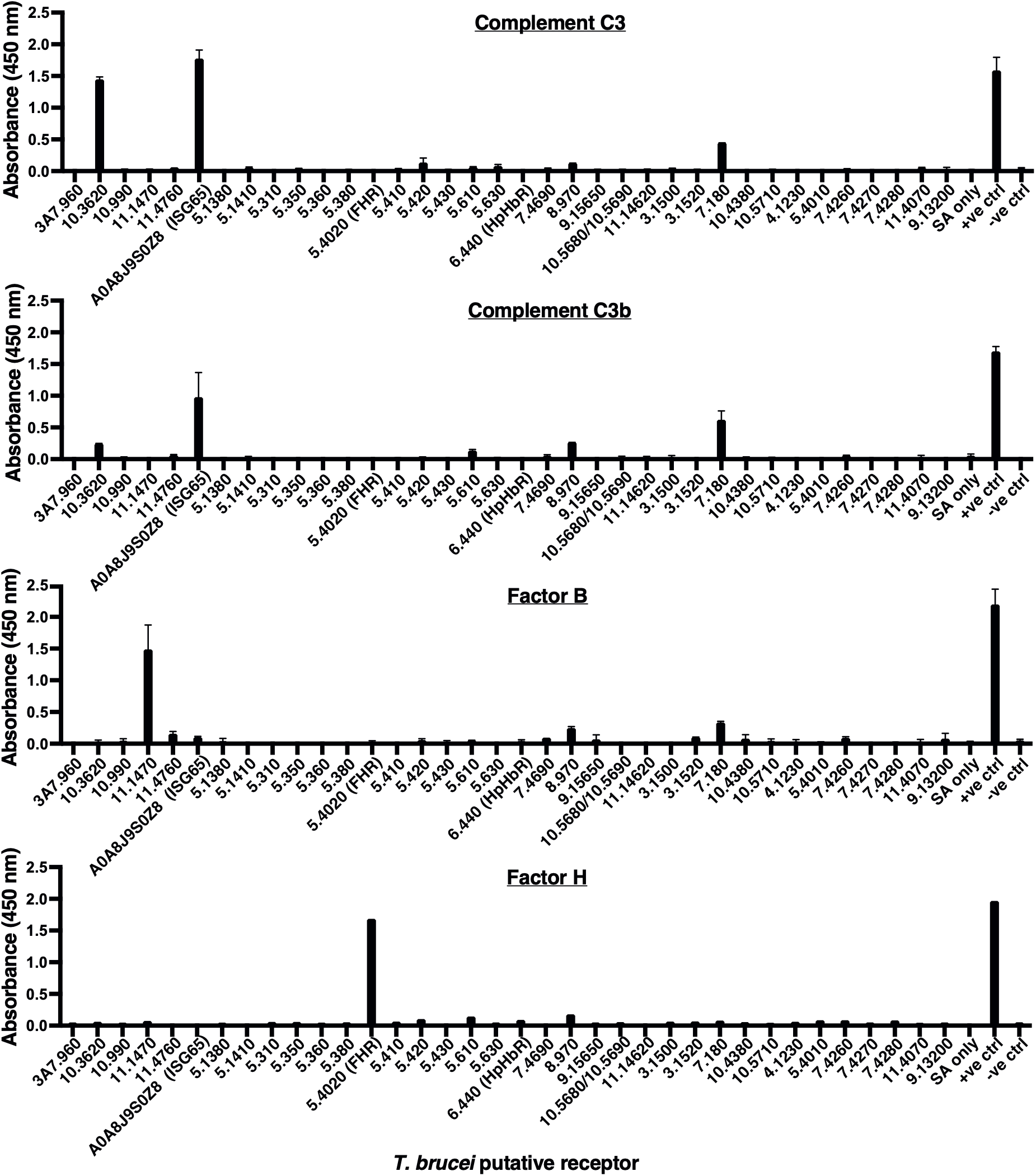
Raw data for complement C3, C3b, Factor B and Factor H binding to the panel of receptors, related to the binding screen in Figure 1c. C3, C3b, and Factor H have known receptors.

**Supplementary Figure 3.**
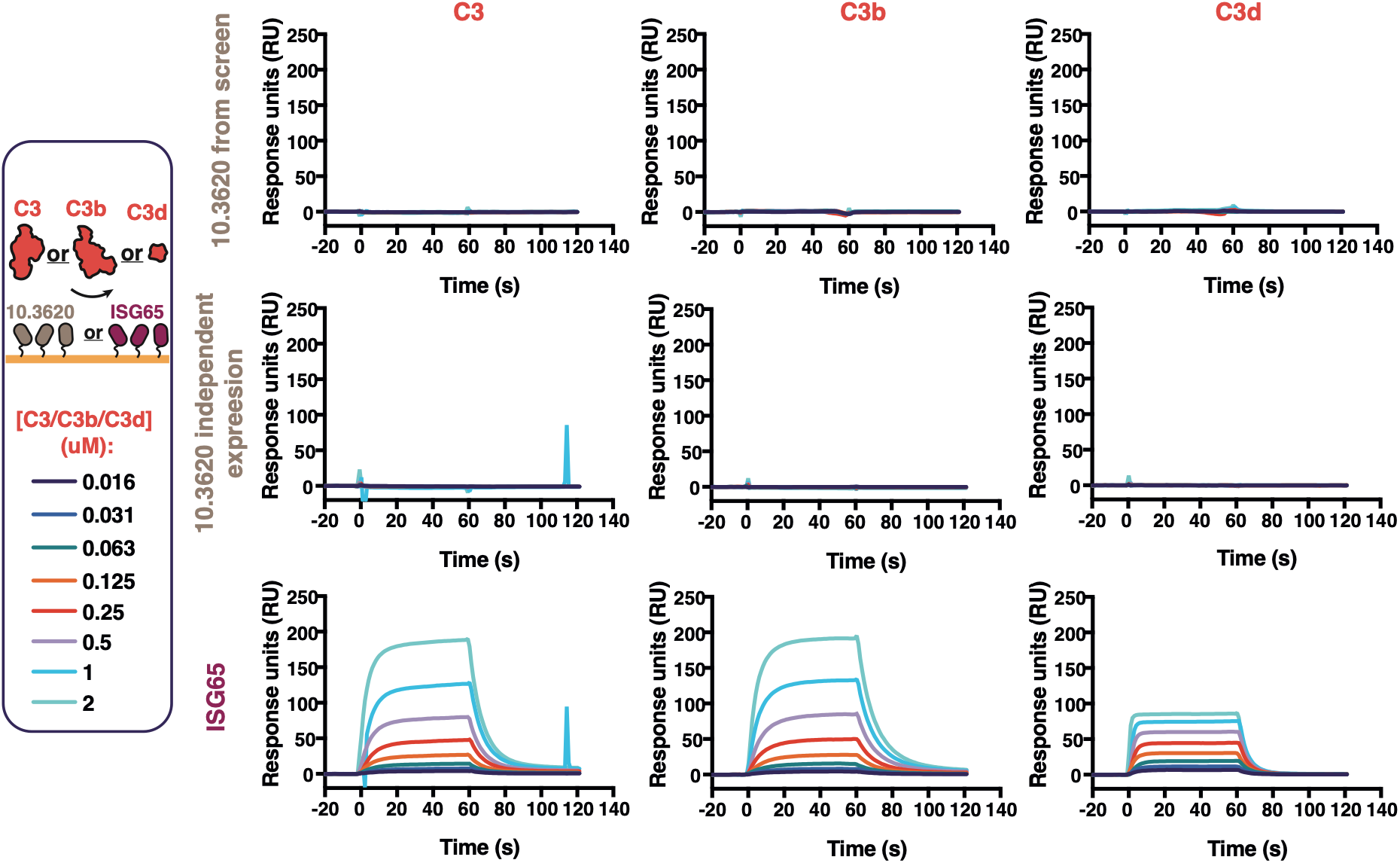
Surface plasmon resonance data showing responses from injection of C3, C3b, or C3d (two-fold serial dilutions from a concentration of 2 μM) over a flow cell with coupled with streptavidin tetramers conjugated to either the batch of Tb927.10.3620 used in the screen in Figure 1c, an independently purified biotinylated Tb927.10.3620, ISG65 as a positive control

**Supplementary Figure 4.**
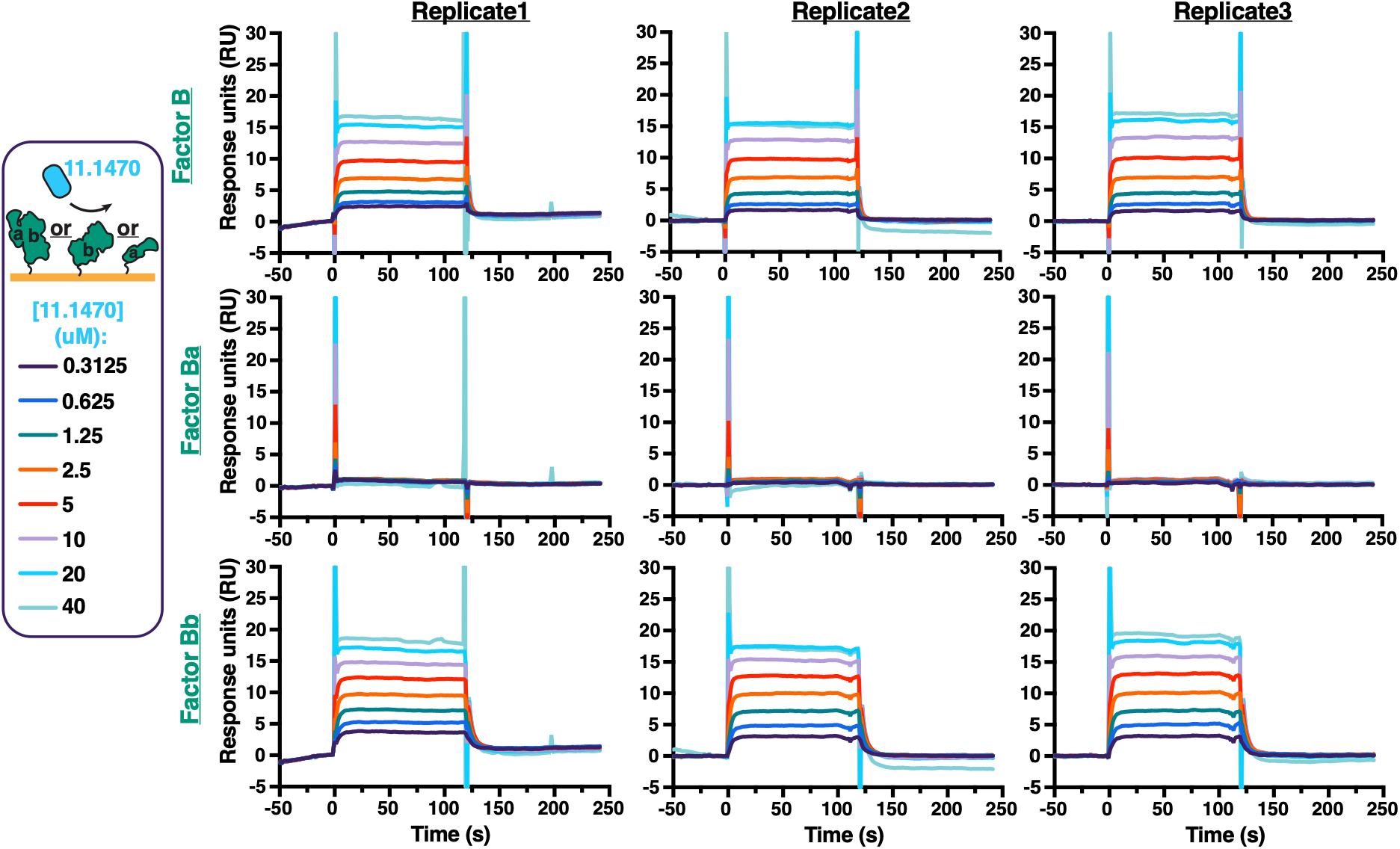
Individual repeats of SPR data in Figure 1d, showing Tb927.11.1470 binding to Factor B, Ba, or Bb.

**Supplementary Figure 5.**
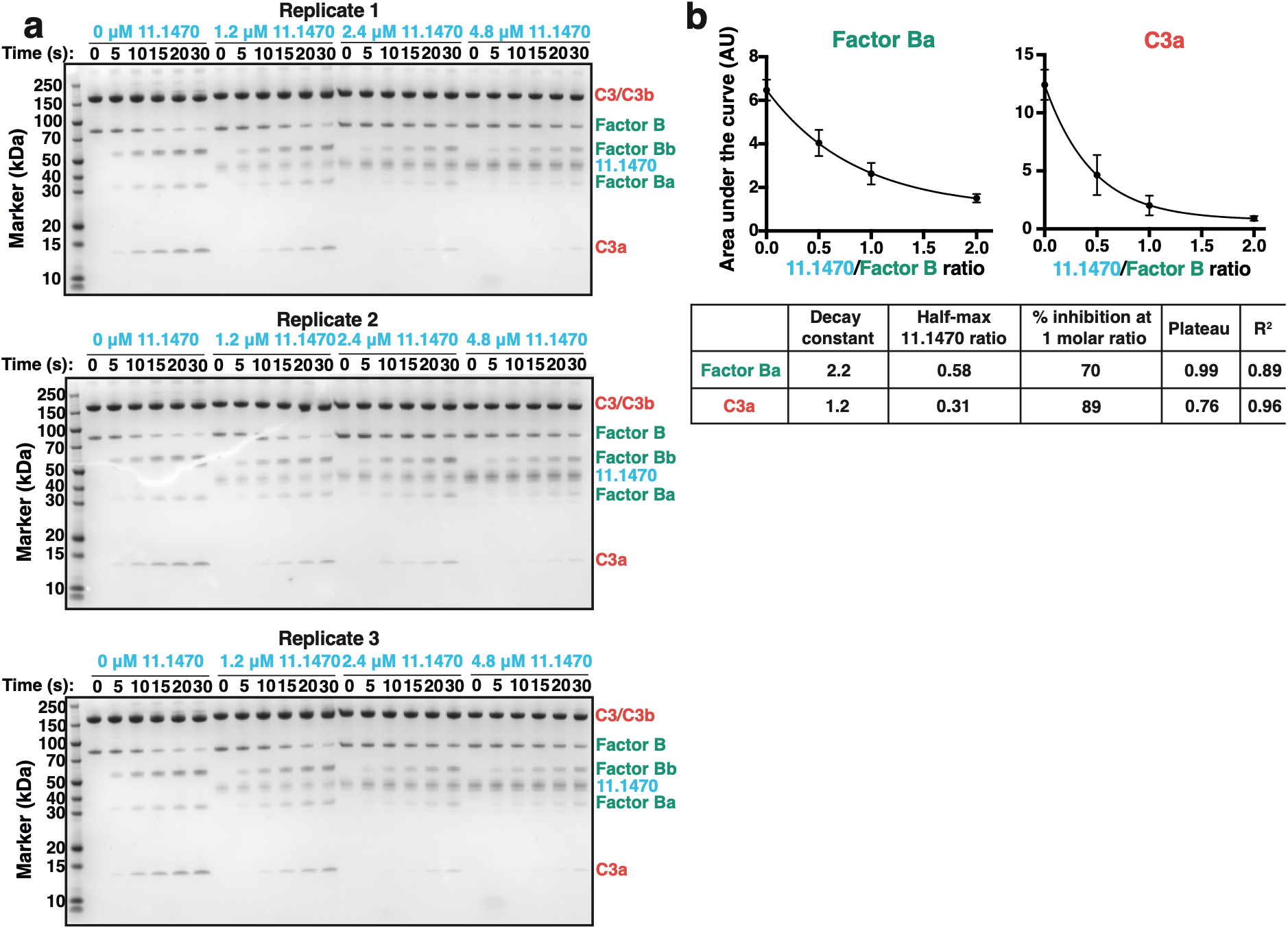
(a) Individual repeats of SDS-PAGE of the C3bBb convertase assay in Figure 2b. (b) Area under the curve analysis of the graphs in Figure 2b, showing how increasing amounts of Tb927.11.1470 decrease production of Factor Ba and C3a over time.

**Supplementary Figure 6.**
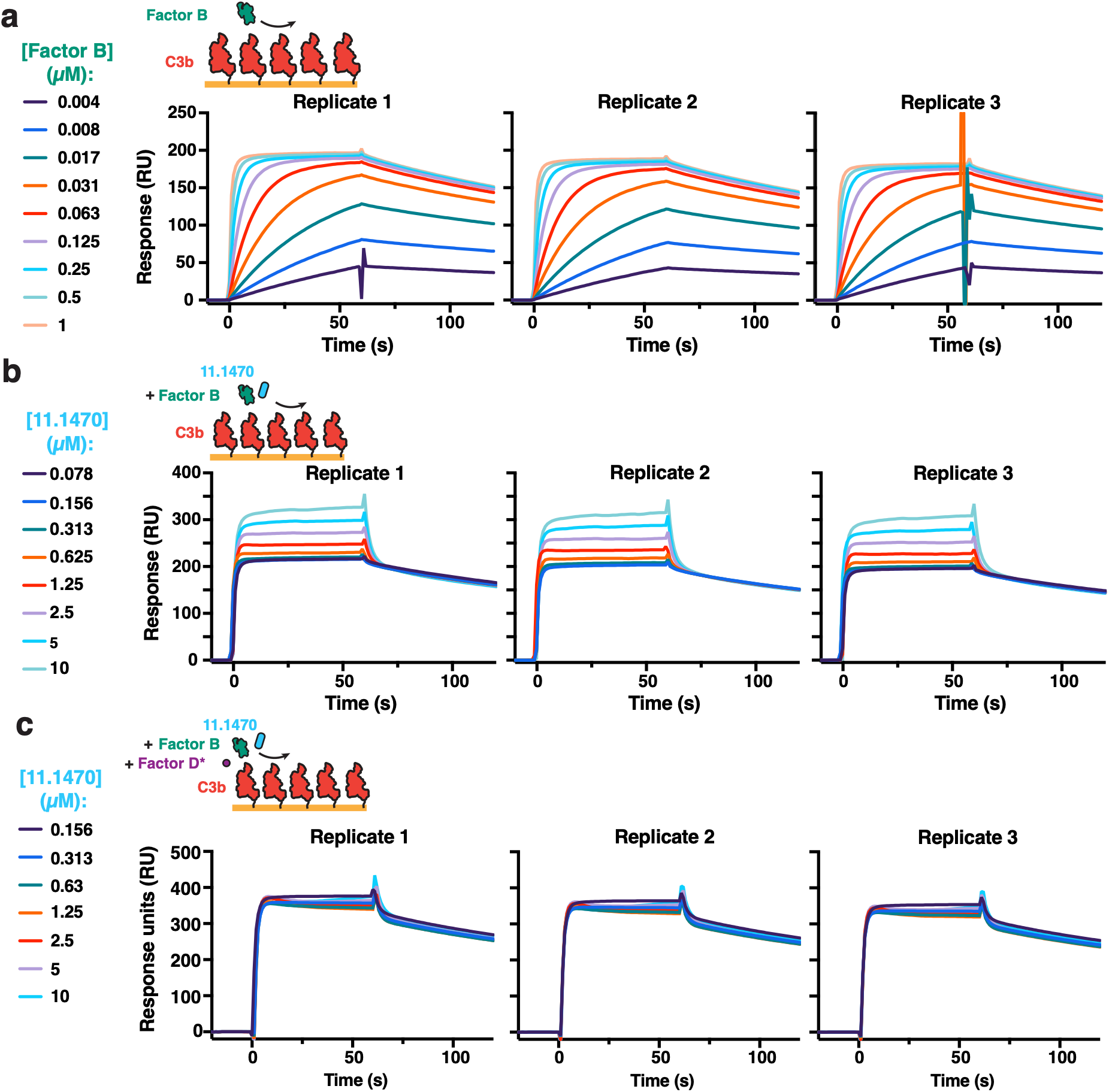
Individual repeats of SPR data in: (a) Figure 3a, showing Factor B binding to C3, (b) Figure 3b, showing Factor B and Tb927.11.1470 binding to C3b, (c) Figure 3c, showing the absence of binding of Tb927.11.1470 to C3b in the presence of Factor B and Factor D.

**Supplementary Figure 7.**
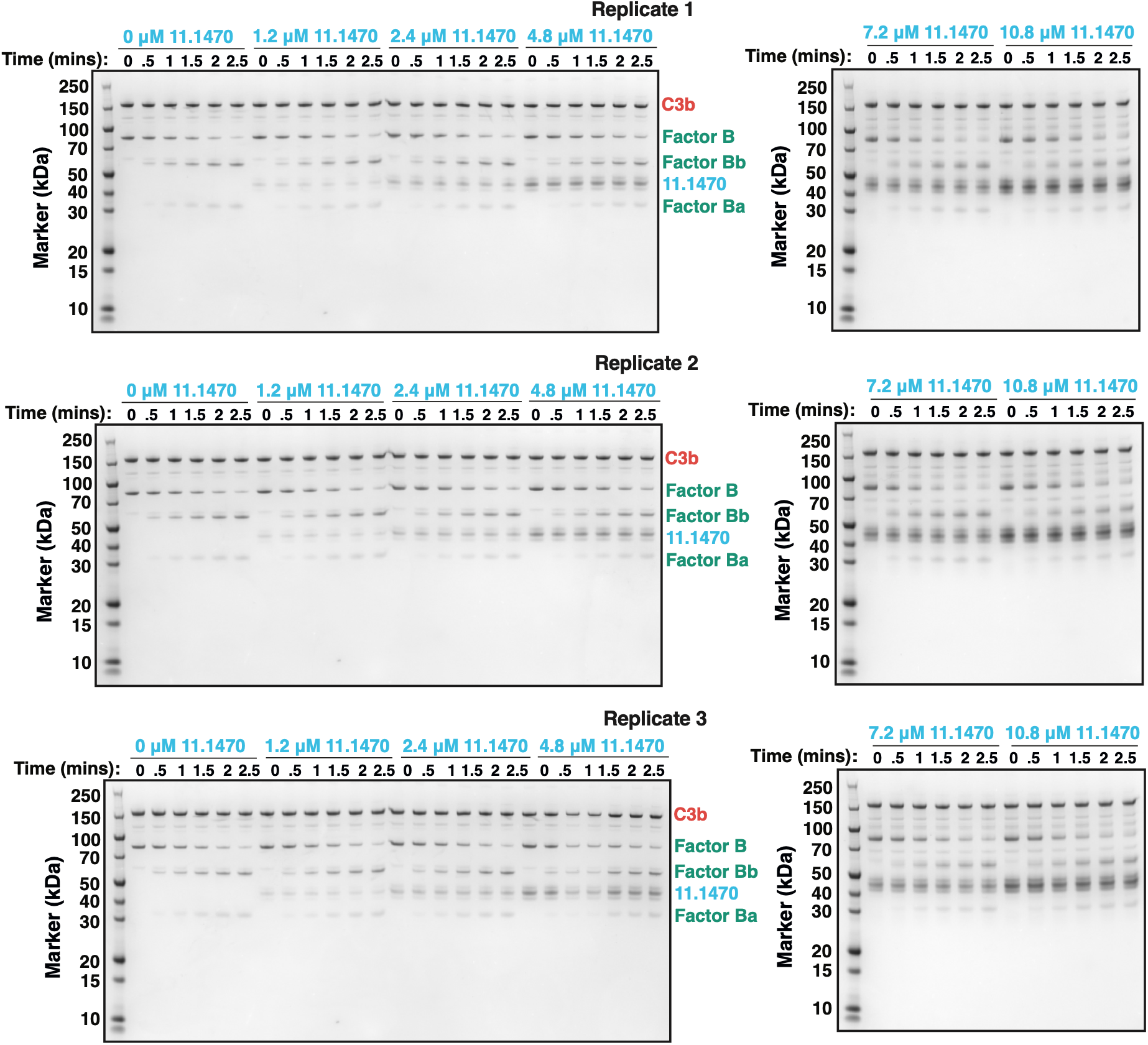
Individual repeats of SDS-PAGE of the C3bB cleavage assay showing in Figure 3d in the presence of increasing amounts of Tb927.11.1470.

**Supplementary Figure 8.**
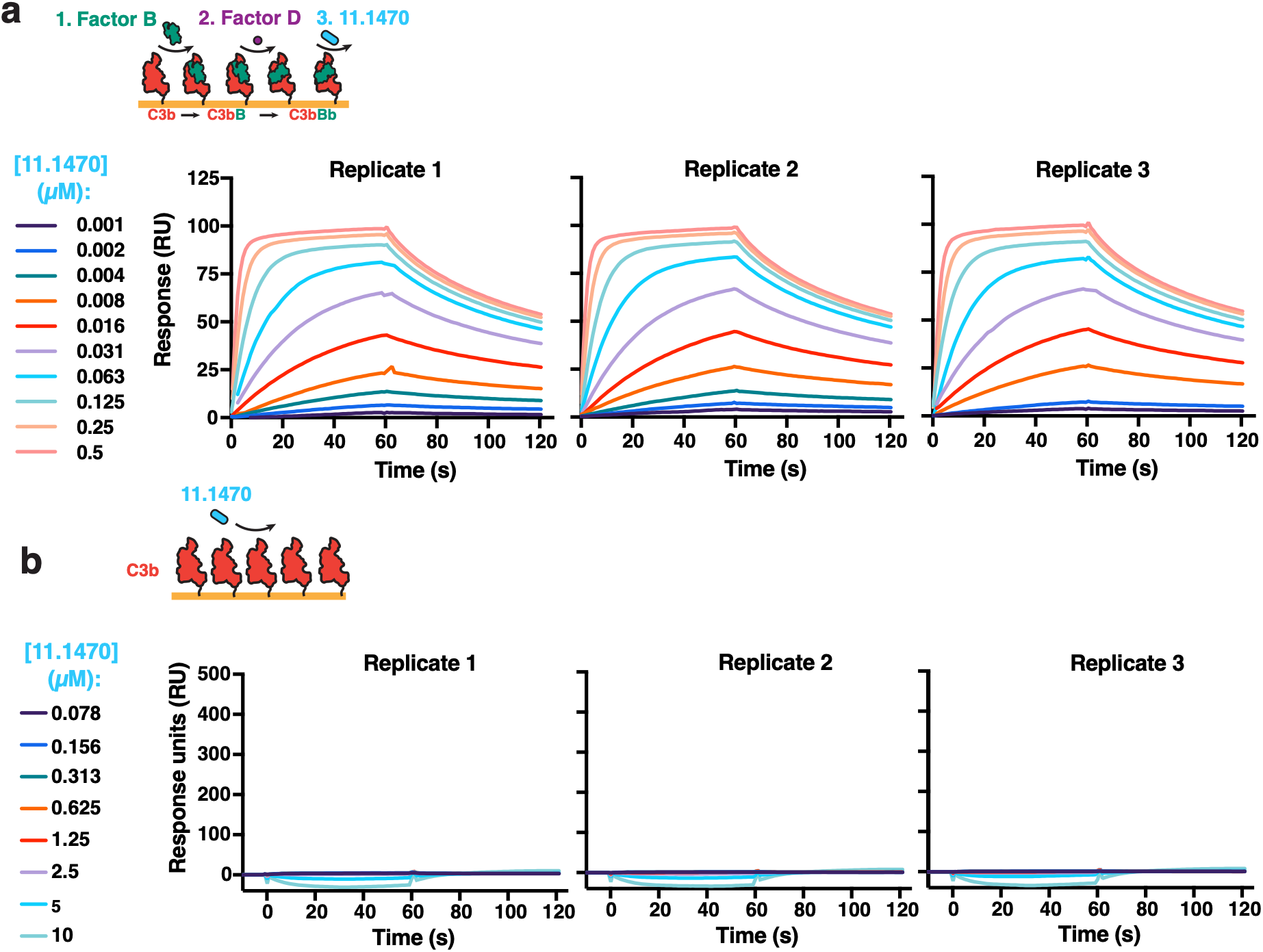
(a) Individual repeats of SPR data in Figure 3e, showing Tb927.11.1470 binding to C3bBb. (b) Individual repeats of SPR data in Figure 3f, showing Tb927.11.1470 binding to C3b.

**Supplementary Figure 9.**
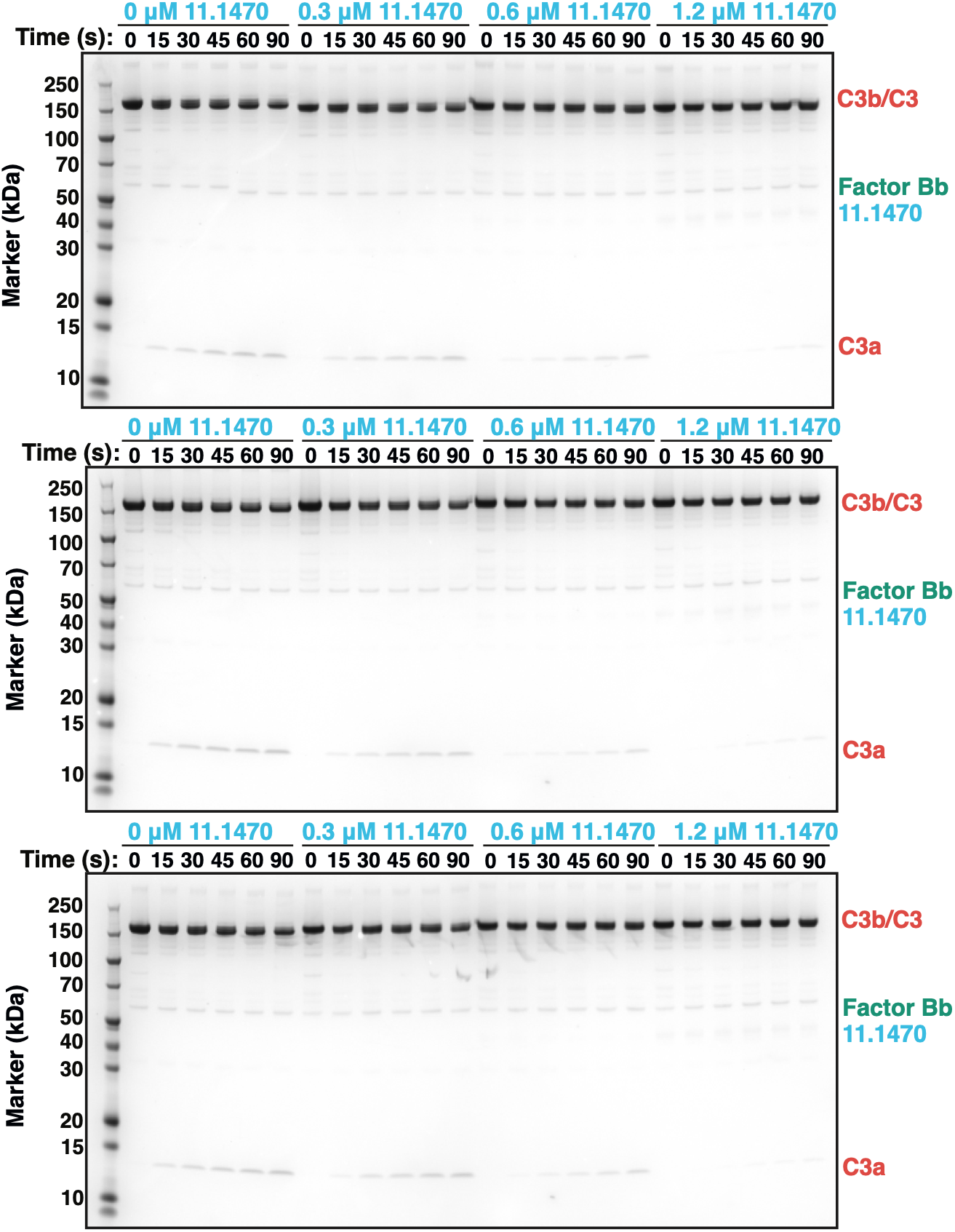
Individual repeats of SDS-PAGE of the C3bBb based C3 cleavage assay shown in Figure 3g, in the presence of increasing amounts of Tb927.11.1470.

**Supplementary Figure 10.**
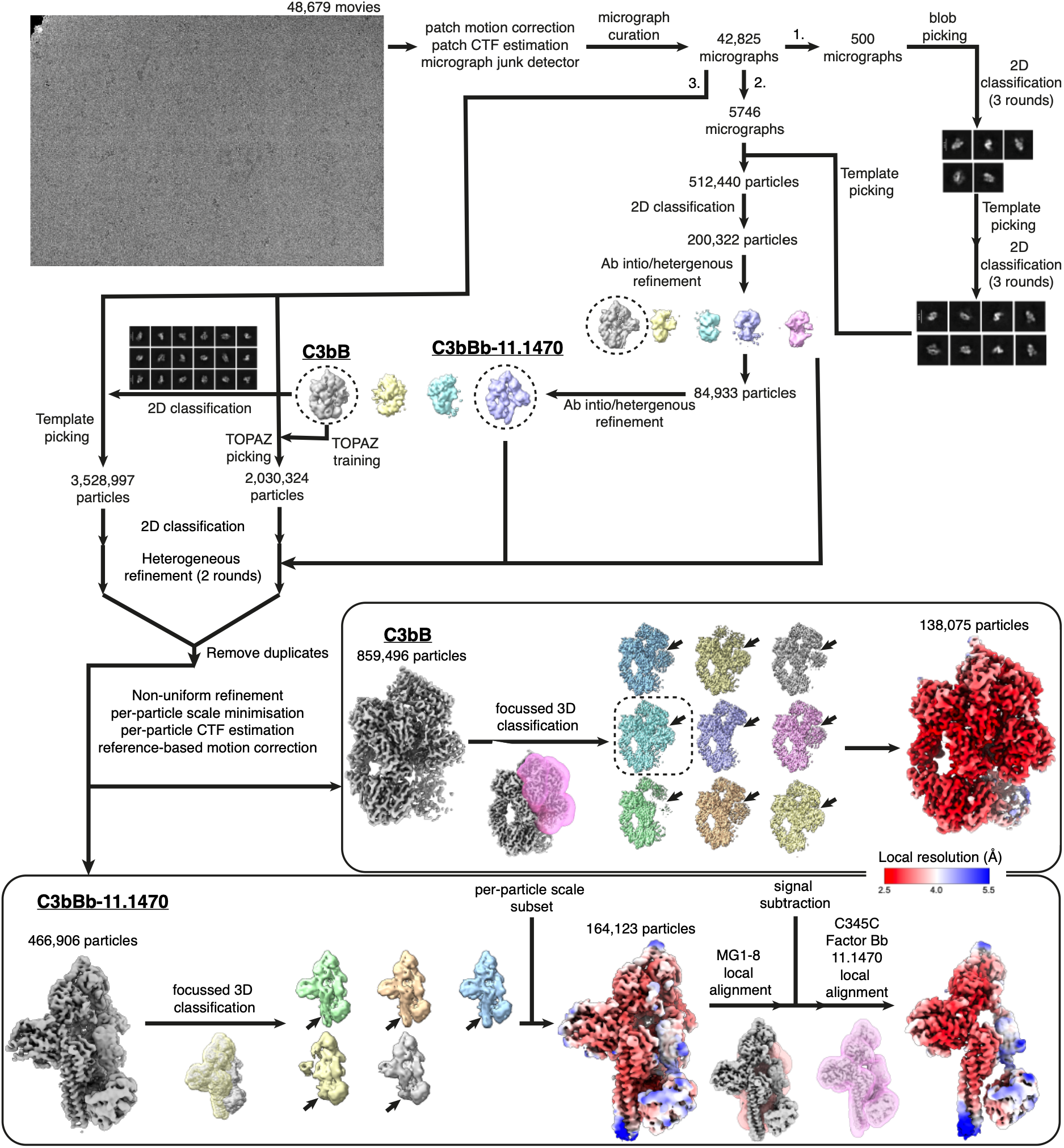
Cryo-EM data processing pipeline of C3bBb bound to Tb927.11.1470. After the arrow labelled 1, initial 2D classes for template picking were generated from 500 micrographs. After arrow 2, a clean stack of particles were obtained from 5746 micrographs, which were used to generate optimal 2D classes for template picking, and to train a TOPAZ model for particle picking. After arrow 3, Template and TOPAZ picks were subjected to 2D classification and heterogeneous refinement, using initial models generated in 2. These particles stacks were merged and duplicate particles removed, generating a particle stack corresponding to uncleaved C3bB, and C3bBb-Tb927.11.1470 complexes. 3D classification improved the occupancy of C3bBb and C3bBb-Tb927.11.1470, with local alignment of C3bBb-Tb927.11.1470 particularly improving the resolution of some regions of Factor Bb.

**Supplementary Figure 11.**
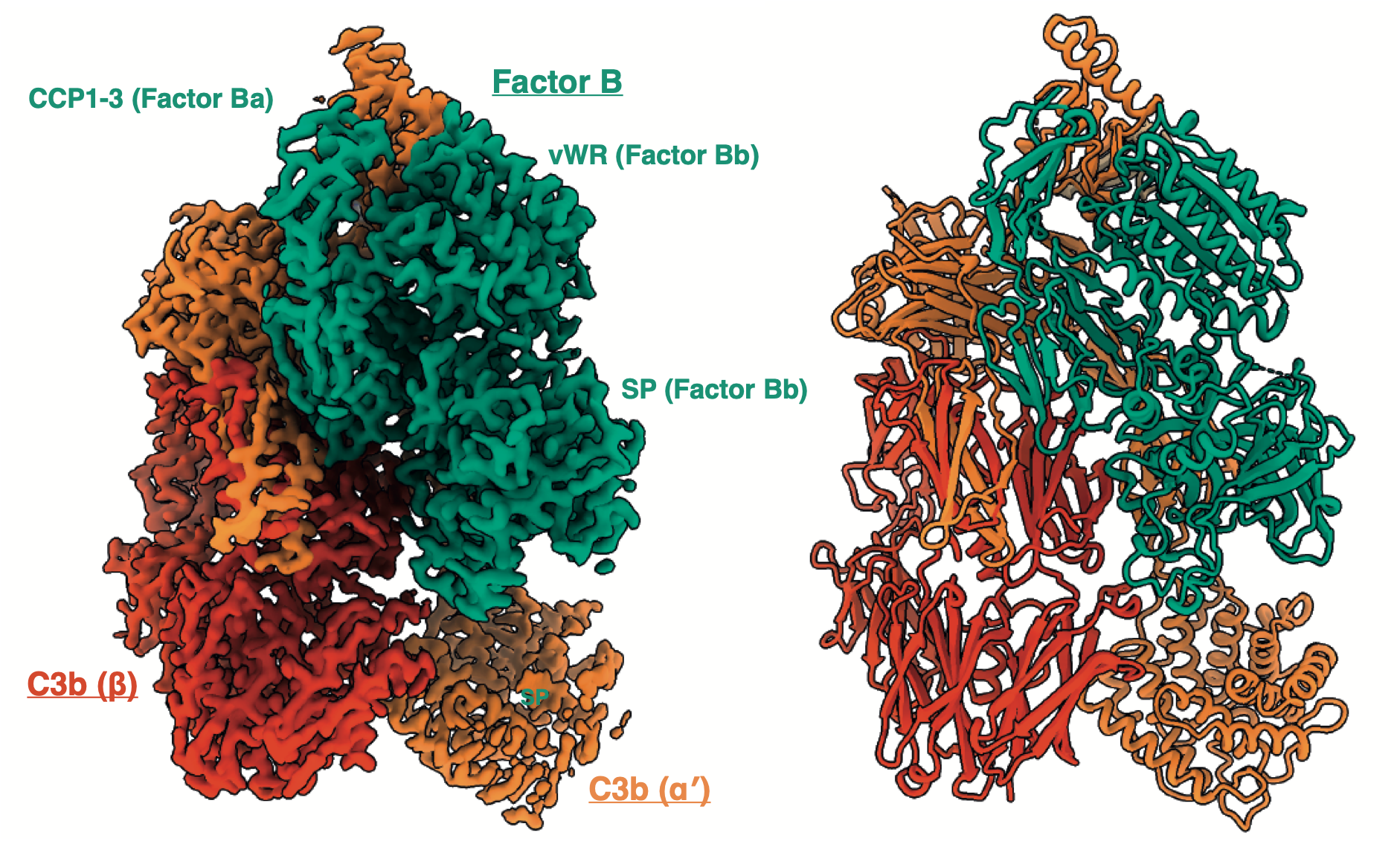
Structure of the proconvertase C3bB at 2.7 Å resolution. Image processing of the cryo-EM dataset of C3bBb bound to C3bBbR revealed some uncleaved C3bB (Supplementary Figure 10).

**Supplementary Figure 12.**
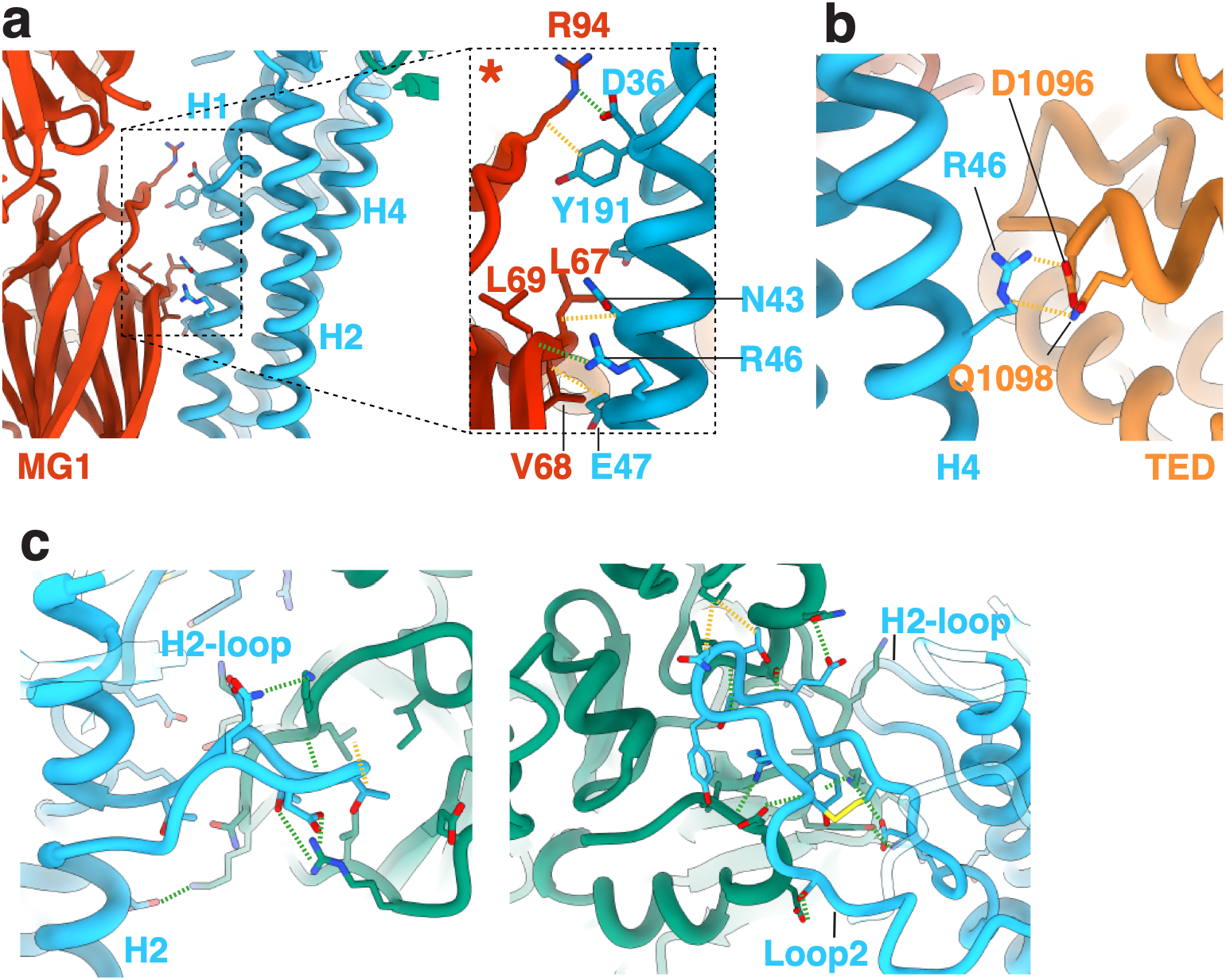
Close up views of the interaction between Tb927.11.1470 and C3bBb. (a) The interaction between Helix 1 (H1) of Tb927.11.1470 and the MG1 domain of the C3b β-chain. (b) The interaction between H2 of Tb927.11.1470 and the thioester domain (TED) of the C3b α-chain. (c) The interaction between Tb927.11.1470 and the Factor Bb serine protease domain showing: on the left, the H2-loop which forms from a break in H2, and on the right, Loop 2 of Tb927.11.1470.

**Supplementary Figure 13.**
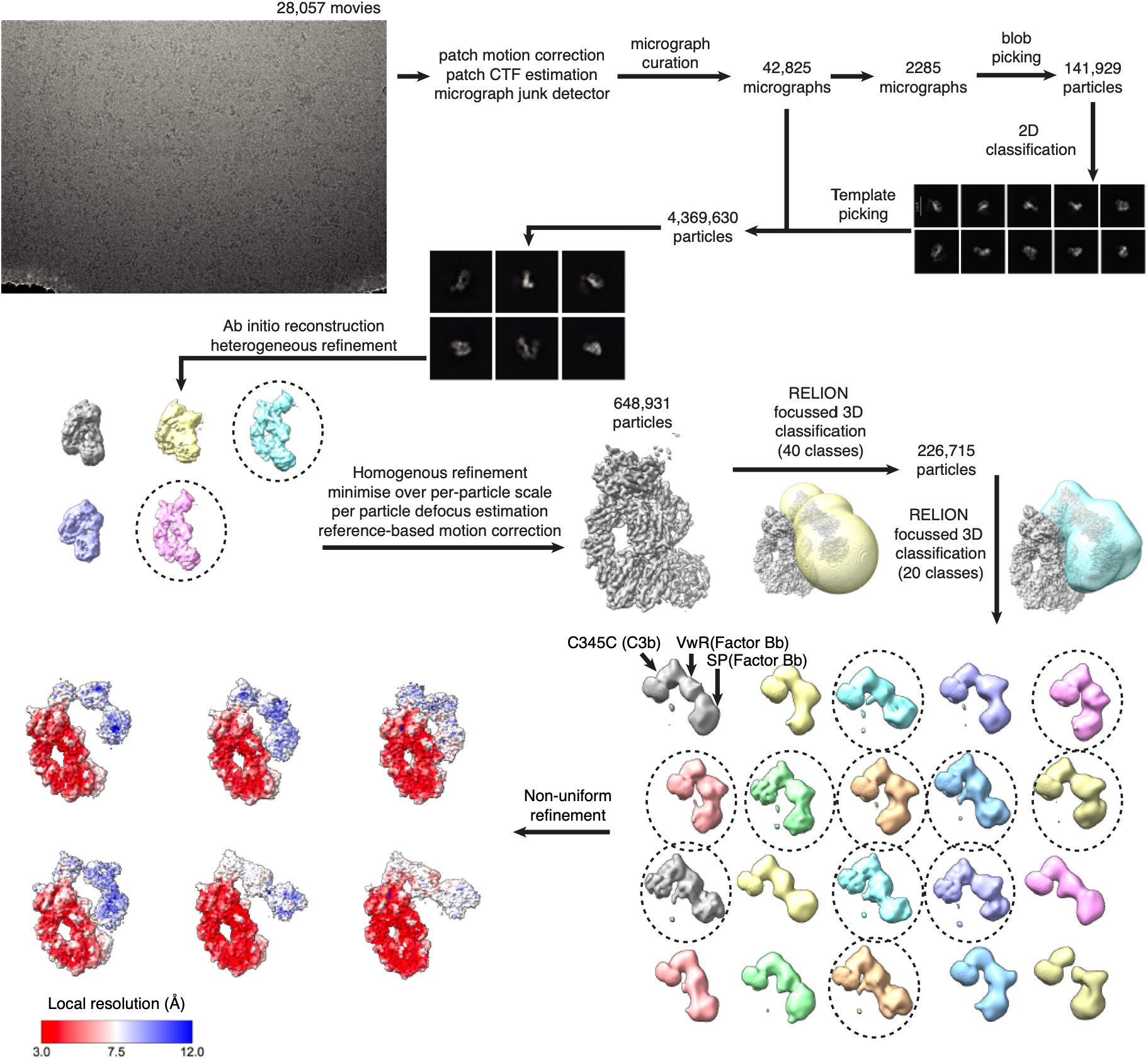
Cryo-EM data processing pipeline for C3bBb. A stack of clean C3bBb particles was obtained from CryoSPARC using 2D classification. *Ab initio* reconstruction and heterogenous refinement. A mask (yellow) encompassing the flexible C345C domain from C3b which is bound to Factor Bb was made from spheres generated in ChimeraX was used to perform focussed 3D classification. A second mask (blue) was generated from all classes containing density for Factor Bb. This was used in a second round of focussed 3D classification, resulting in 20 classes (displayed in various colours). Classes with dotted lines were selected for non-uniform refinement in CryoSPARC.

## Supplementary Tables

**Supplementary Table 1.** Putative *T. brucei brucei* cell surface receptors identified in the bioinformatic screen. Protein names are listed under ‘Protein’. For gene families, representative members were selected. For recombinant protein production, the start and end amino acid included in expression plasmids is noted under ‘Construct boundaries’, with the final cell line and culture volume used for expression noted under ‘Expression’. *For consistency with previous publications^22,23^ a representative ISG65 sequence from the

| Protein | Protein Length | Construct boundaries | Expression |
| --- | --- | --- | --- |
| Tb06.3A7.960 | 310 | 23-287 | S2 (50 mL) |
| Tb927.10.3620 | 308 | 19-285 | HEK293F (250 mL) |
| Tb927.10.990 | 384 | 25-356 | HEK293F (50 mL) |
| Tb927.11.1470 | 437 | 24-385 | HEK293F (50 mL) |
| Tb927.11.4760 | 272 | 22-249 | S2 (700 mL) |
| A0A8J9S0Z8 (ISG65)* | 436 | 24-385 | HEK293F (50 mL) |
| Tb927.5.1380 | 285 | 20-266 | S2 (50 mL) |
| Tb927.5.1410 | 435 | 22-376 | HEK293F (50 mL) |
| Tb927.5.310 | 407 | 22-346 | HEK293F (50 mL) |
| Tb927.5.350 | 522 | 29-467 | HEK293F (50 mL) |
| Tb927.5.360 | 523 | 29-468 | HEK293F (50 mL) |
| Tb927.5.380 | 552 | 27-481 | S2 (50 mL) |
| Tb927.5.4020 (FHR) | 272 | 25-252 | HEK293F (50 mL) |
| Tb927.5.410 | 393 | 29-342 | HEK293F (50 mL) |
| Tb927.5.420 | 373 | 24-325 | HEK293F (50 mL) |
| Tb927.5.430 | 377 | 24-333 | HEK293F (50 mL) |
| Tb927.5.610 | 428 | 23-373 | HEK293F (50 mL) |
| Tb927.5.630 | 401 | 25-349 | HEK293F (50 mL) |
| Tb927.6.440 (HpHbR) | 403 | 33-375 | HEK293F (50 mL) |
| Tb927.7.4690 | 319 | 20-267 | S2 (50 mL) |
| Tb927.8.970 | 459 | 21-438 | S2 (50 mL) |
| Tb927.9.15650 | 346 | 25-324 | S2 (50 mL) |
| Tb927.10.5680/90 | 394 | 21-374 | HEK293F (500 mL) |
| Tb927.11.14620 | 458 | 18-434 | HEK293F (50 mL) |
| Tb927.3.1500 | 366 | 22-366 | HEK293F (250 mL) |
| Tb927.3.1520 | 365 | 19-346 | HEK293F (50 mL) |
| Tb927.7.180 | 437 | 22-437 | S2 (50 mL) |
| Tb927.10.4380 | 246 | 21-230 | S2 (50 mL) |
| Tb927.10.5710 | 227 | 23-208 | HEK293F (50 mL) |
| Tb927.4.1230 | 331 | 28-305 | HEK293F (50 mL) |
| Tb927.5.4010 | 304 | 19-278 | HEK293F (50 mL) |
| Tb927.7.4260 | 263 | 25-189 | S2 (50 mL) |
| Tb927.7.4270 | 222 | 29-169 | S2 (50 mL) |
| Tb927.7.4280 | 211 | 28-176 | S2 (700 mL) |
| Tb927.11.4070 | 224 | 20-201 | HEK293F (50 mL) |
| Tb927.9.13200 | 189 | 17-172 | S2 (50 mL) |

**Supplementary Table 2.**
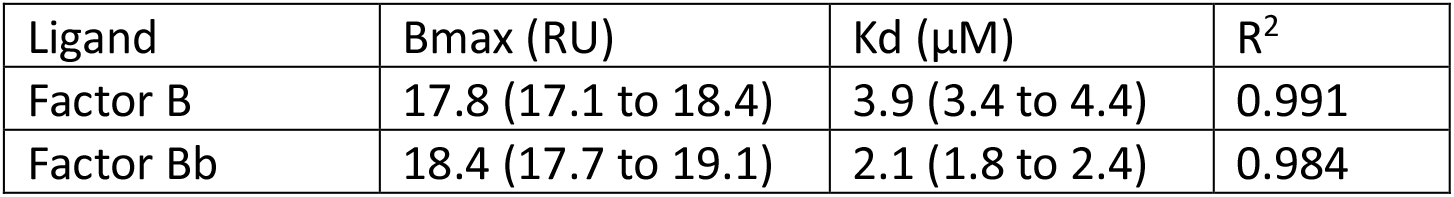
Steady state affinity parameters for 11.1470 binding to Factor B and Factor Bb. 95 % confidence intervals are indicated in brackets; n = 3 experimental replicates.

**Supplementary Table 3.** Steady state affinity parameters for Factor B binding to C3b. 95 % confidence intervals are indicated in brackets; n = 3 experimental replicates.

| Ligand | Bmax (RU) | Kd (μM) | R<sup>2</sup> |
| --- | --- | --- | --- |
| C3b | 196 (191 to 202) | 0.011 (0.010 to 0.013) | 0.974 |

**Supplementary Table 4.** Steady state affinity parameters for 11.1470 binding to C3bB. 95 % confidence intervals are indicated in brackets; n = 3 experimental replicates.

| Ligand | Bmax (RU) | Kd (μM) | Background | R<sup>2</sup> |
| --- | --- | --- | --- | --- |
| C3bB | 168 (140 to 215) | 4.5 (2.7 to 8.2) | 200 (193 to 207) | 0.957 |

**Supplementary Table 5.** Steady state affinity parameters for 11.1470 binding to C3bBb. 95 % confidence intervals are indicated in brackets; n = 3 experimental replicates.

| Ligand | Bmax (RU) | Kd (μM) | R<sup>2</sup> |
| --- | --- | --- | --- |
| C3bBb | 106 (101 to 111) | 0.023 (0.020 to 0.028) | 0.996 |

**Supplementary Table 6.** Kinetic parameters for 11.1470 binding to C3bBb. Standard deviation is indicated; n = 3 experimental replicates.

| Ligand | k<sub>a</sub> (M<sup>-1</sup>s<sup>-1</sup>) | k<sub>d</sub> (s<sup>-1</sup>) | K<sub>D</sub> (μM) | Rmax | Chi<sup>2</sup> |
| --- | --- | --- | --- | --- | --- |
| C3bBb | 1.11x10<sup>6</sup> | 1.07x10<sup>-2</sup> | 9.7 ± 0.5 | 95 ± 1.1 | 0.6 ± 0.2 |

**Supplementary Table 7.**
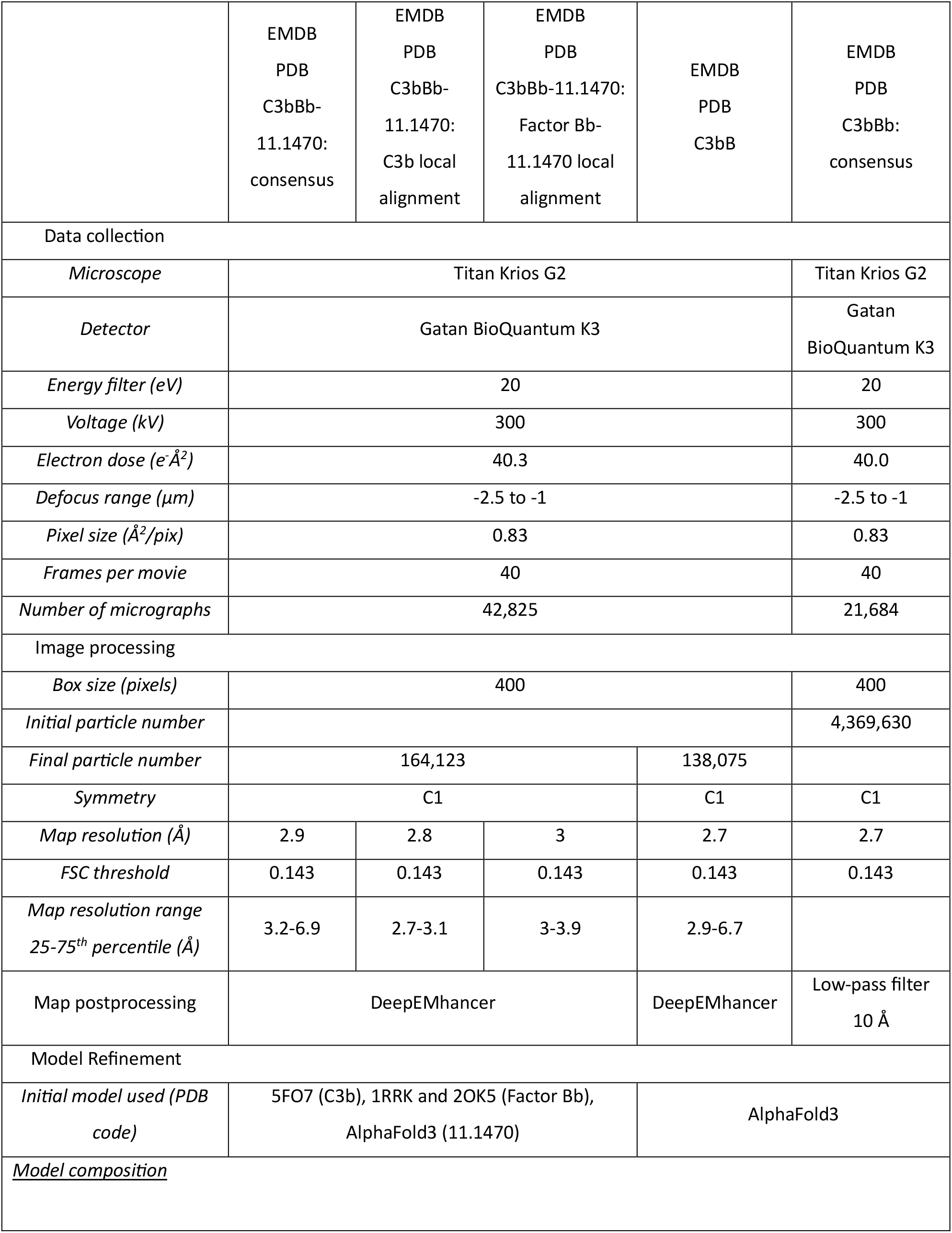
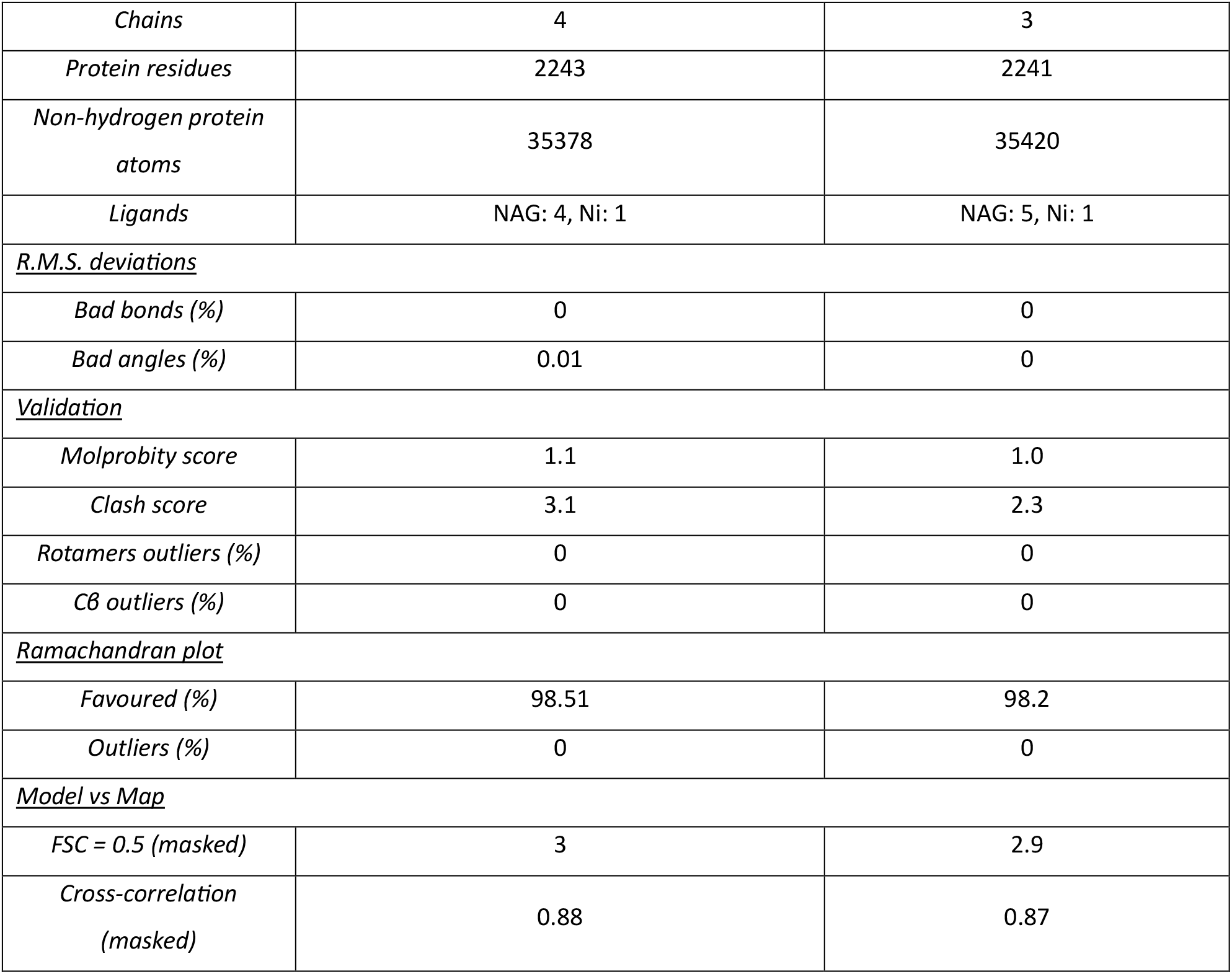
Cryo-EM data collection and refinement statistics. Statistics for Model composition, R.M.S deviations, validation and Ramachandran plot were calculated in MolProbity^63^, Statistics for Model vs Map were calculate in Phenix.

